# Lipid-conjugated DNA enables on-demand delivery of lipids and proteins to synthetic cells

**DOI:** 10.64898/2026.03.30.715215

**Authors:** Bert Van Herck, Jacob Kerssemakers, Nikolaj A. Risgaard, Stefan Vogel, Cees Dekker, Gijsje H. Koenderink

## Abstract

The bottom-up construction of synthetic cells based on giant unilamellar vesicles (GUVs) is a central goal in synthetic biology. Achieving targeted changes in membrane and cytoplasmic composition with temporal control remains challenging however. DNA-mediated fusion with small vesicles (∼100 nm large unilamellar vesicles; LUVs) has been proposed as a strategy to deliver lipids and cytosolic contents in a programmable manner. However, *in vitro*, membrane fusion is generally found to be inefficient and poorly controllable for reasons that are poorly understood. Here, we present an approach based on lipid-conjugated DNA (LiNA) to mediate programmable fusion between LUVs and micron-sized GUVs, which we quantitatively monitor with confocal microscopy at the single-GUV level. We show that lipid and content mixing both occur with high efficiency over a wide range of LiNA concentrations, demonstrating that LiNAs indeed induce robust membrane fusion. Furthermore, we show that LiNA-mediated fusion provides a powerful tool to deliver cytosolic biomolecules, enabling control over internal activities. Our findings establish a quantitative framework for studying fusion-driven processes in synthetic cells and provide a versatile platform for the programmable delivery of lipids and cytosolic cargoes - thus advancing the development of synthetic cells that can grow and adapt through fusion-based uptake of molecular building blocks.

## Introduction

Cellular life emerged from simple molecular assemblies whereupon billions of years of evolution shaped the cellular complexity and diversity that we encounter today. Cells represent the smallest self-replicating structural and functional units of life, maintaining homeostasis within a compartment that is confined by a lipid-bilayer membrane. This compartment is maintained out of thermodynamic equilibrium, enabling metabolic reactions and proliferation^1–3^. Recently, efforts have been starting to recreate life-like behaviors in synthetic cell-like systems from the bottom-up, promising a new route towards understanding the fundamental mechanisms governing biological organization and function^4^.

Giant unilamellar vesicles (GUVs) are commonly used as cell mimics as their supramolecular architecture - a phospholipid bilayer surrounding a micron-sized aqueous compartment - resembles that of a cell^5^. Although phospholipid membranes are permeable to small hydrophobic and neutral molecules, they are impermeable to most hydrophilic molecules (including proteins) as their transport involves the crossing of the hydrophobic membrane core. Consequently, GUVs behave as effectively closed systems in which all building blocks must be encapsulated from the onset^6^. This represents a significant limitation for the bottom-up reconstitution of synthetic cells as some cellular components require replenishment over time and reactions may need to be initiated in a controlled and programmable manner after GUV formation.

Notably, unlike cells, simple GUV-like synthetic cells lack any specialized membrane protein transporters. GUVs can be rendered permeable through the insertion of pore-forming toxins (e.g. *α*-haemolysin^7^) or artificial nanopores engineered with DNA origami^8–10^ in the membrane. However, such pores have limited selectivity, being selective only for cargo size. Also, they are non-directional and unable to prevent encapsulated molecules from diffusing out, and they cannot drive the accumulation of solutes against their concentration gradient^3^. The *in vitro* reconstitution of transmembrane proteins that selectively transport biomolecules across the membrane remains challenging, requiring complex reconstitution procedures and suffering low reconstitution efficiencies^11^. As a result, the controlled and timed delivery of biomolecules into GUVs remains challenging, highlighting the need for an approach to precisely alter the composition of synthetic cells on demand.

Delivering pre-synthesized molecules or precursors directly to synthetic cells through membrane fusion with nanoscale carrier vesicles provides an appealing and efficient route to support membrane expansion and intracellular enrichment, conceptually resembling the process of endocytosis^12^ in cells. Central to this approach is the fusion of lipid membranes, a ubiquitous and highly regulated process that is essential for living cells as it underlies organelle fusion^13^, endosomal trafficking^14^, viral infection^15^, and neurotransmission^16,17^. Membrane fusion occurs through a series of carefully orchestrated stages^18,19^: Initially, the membranes of two opposing vesicles are brought into close contact (docking), which sets the stage for membrane destabilization, culminating in the formation of a hemifusion diaphragm. In this state, the outer leaflets of the bilayers have fused (and lipids can mix between them), but the inner leaflets and the aqueous compartments remain separate. Subsequently, a small fusion pore opens within the hemifusion diaphragm, initially flickering between open and closed states. With time, this pore stabilizes and expands, permitting the mixing of both the lipids and the aqueous contents between the fusing compartments. This final stage marks the complete merger of the two membranes, achieving full membrane fusion.

Membrane fusion is controlled by specialized membrane fusion protein machineries, amongst which the Soluble NSF Attachment Protein Receptor (SNARE) protein family is studied the most extensively^16,20^. These proteins are highly conserved among eukaryotes and homologous genes have been identified in Asgard archaea and *γ*-proteobacteria^21^. These fusogens play a crucial role in overcoming the repulsive hydration barrier, facilitating direct van der Waals interactions between the lipid headgroups and bringing the membranes closer together, thereby setting the stage for membrane fusion. Reconstituted SNAREs have been used to study membrane fusion *in vitro*^22–26^. Nevertheless, the use of SNAREs recquires meticulous protein purification, correct orientation in the membrane, and additional regulatory proteins (belonging to the Sec1/Munc18-like or CATCHR protein families) in order to establish efficient membrane fusion. As a result of this complexity, the exact assembly pathway and dynamics of SNARE complex assembly are still not completely understood.

To reduce complexity, approaches have been developed that mimic SNAREs, including lipid-conjugated nucleic acids (LiNAs). These tagged DNA molecules have been shown to insert spontaneously into membranes^27–29^ and induce fusion between distinct populations of large unilamellar vesicles (LUVs)^30–32^, as well as between LUVs and supported lipid bilayers^33–35^. The concept where LiNAs with a complementary sequence are used on both vesicles does in principle allow to ‘program’ certain classes of LUVs that can subsequently target the fusion to other defined vesicles by hybridization, allowing on-demand delivery of constituents.

However, previous DNA-based approaches using one or two terminal membrane anchors were hampered by either incomplete fusion, characterized by only moderate levels of inner leaflet mixing and content mixing, or substantial membrane leakage^30–32,35^. To overcome these limitations, different anchoring strategies have been developed that employ membrane-spanning anchors^35^ or lipid-conjugated nucleotides^36–38^.

The *in vitro* characterization of membrane fusion remains difficult due to the dynamic character of the process and the inability to directly observe the nanoscale intermediate states with optical methods. Classical bulk fluorescence assays, such as lipid mixing or content mixing between populations of vesicles, provide only ensemble averages^39–42^. Because the signal arises from the average of a large and heterogeneous vesicle population, critical information about individual fusion events is lost. Often, changes in fluorescence intensity cannot be unambiguously attributed to membrane fusion, as similar readouts may arise from vesicle leakage, rupture, or photobleaching. Moreover, variations in vesicle size, lamellarity, or encapsulation efficiency further complicate the interpretation of bulk data^43^.

In contrast, single-vesicle measurements on micron-sized GUVs offer a more direct and reliable assessment of membrane fusion. Individual vesicles can be monitored over time, allowing clear distinction between fusion, leakage, and rupture, and revealing the population heterogeneity in fusion efficiency and kinetics that is inaccessible in bulk assays^43^. Furthermore, the measurement of GUV diameter and morphology provides an additional layer of information. Indeed, the use of single GUVs for studying membrane fusion is gaining more traction and has been applied to study varying fusogens, including peptides and proteins^41,44–47^ and oppositely charged lipids in the LUV and GUV membranes^48–51^. Yet, *in vitro* membrane fusion remains generally poorly characterized and often inefficient or inconsistent for reasons that are poorly understood.

Here, we employ LiNA fusogens to efficiently mediate the on-demand fusion between LUVs and GUVs. We present two complementary assays to independently quantify lipid and content mixing at the single-GUV level by the combination of fluorescence lifetime and confocal microscopy. By translating the degree of lipid and content mixing into a fusion efficiency metric, we quantify the fusion rate and show that on average a GUV with a radius of 10 *µ*m undergoes 280 fusion events. Real time imaging of the fusion process reveals that full fusion is reached within one hour. Interestingly, we observed a strong temperature dependence which can be used to trigger the (de)activation of fusion^38^. Finally, we show the utility of this fusion system for the programmable delivery of proteins to GUVs. As a proof of concept, we deliver the 55 kDa protein fascin to actin-containg GUVs^52^, triggering on-demand actin bundle formation inside the GUV lumen. These results demonstrate the capability of LiNA-mediated LUV-to-GUV fusion for the delivery of lipids and cytosolic biomolecules into enclosed membrane compartments in a controlled manner, establishing it as a versatile tool for synthetic biology applications.

## Results and discussion

### Fusion system design

The goal of our work was to achieve efficient and controllable fusion between LUVs and GUVs for the on-demand delivery of distinct cargoes, including lipids, small organic molecules, and proteins. We opted for hybridization of complementary DNA molecules on the LUV and GUV surface because of the specific and programmable nature of base pairing. While some studies have employed cholesterol-modified DNA strands^32,53,54^, in our hands this approach was highly inefficient (Supplementary Figure 1, see also prior work^38^). We therefore instead employ DNA constructs with a different membrane anchor that were previously reported to successfully mediate fusion between distinct populations of LUVs^37,38,55^. The construct consists of a 17 nt ssDNA oligonucleotide that is linked via a triethylene glycol spacer to a hydrophobic moiety that spontaneously inserts in lipid membranes to provide stable anchoring (Figure 1A). The anchor structure contains two C16 alkyl chains, as it was demonstrated earlier that a bivalent lipid modification is necessary for irreversible anchoring^56^. Attachment of the anchors to the opposite ends (i.e., to the 5’ and 3’ end) of the complementary oligonucleotides in the LUV and GUV, respectively, results in a zipper configuration upon hybridization (Figure 1B-E). This geometry is reminiscent of SNARE-mediated membrane fusion, where progressive zippering together of the external domains draws the opposing membranes into close proximity^24,57^. Unlike SNARE proteins, these LiNA constructs do not possess intrinsic membrane destabilizing abilities, but are thought to induce spontaneous membrane fusion by tethering the opposing membranes in close proximity, thereby promoting membrane contact through thermal fluctuations^37,42^. In this configuration, the inter-membrane distance is substantially smaller than the common membrane separation of *>* 3 nm that is maintained by hydration and electrostatic repulsion when vesicles exhibit random Brownian motion in solution. Bringing membranes into this sub-3 nm regime sharply increases the energetic penalty associated with the repulsive hydration barrier. Zippering of the complementary LiNA strands expels interfacial water and establishes local adhesion, thereby lowering this barrier and facilitating closer contact. The cumulative effect of multiple duplexes in a membrane–membrane contact zone can become sufficient to induce the local curvature and mechanical stress required for stalk formation^32,42^. This compression of the intermembrane space also imposes entropic penalties on confined lipids and water molecules, further destabilizing the lamellar arrangement and biasing the system towards fusion-pore nucleation and subsequent membrane merging^32,58^.

**Figure 1:**
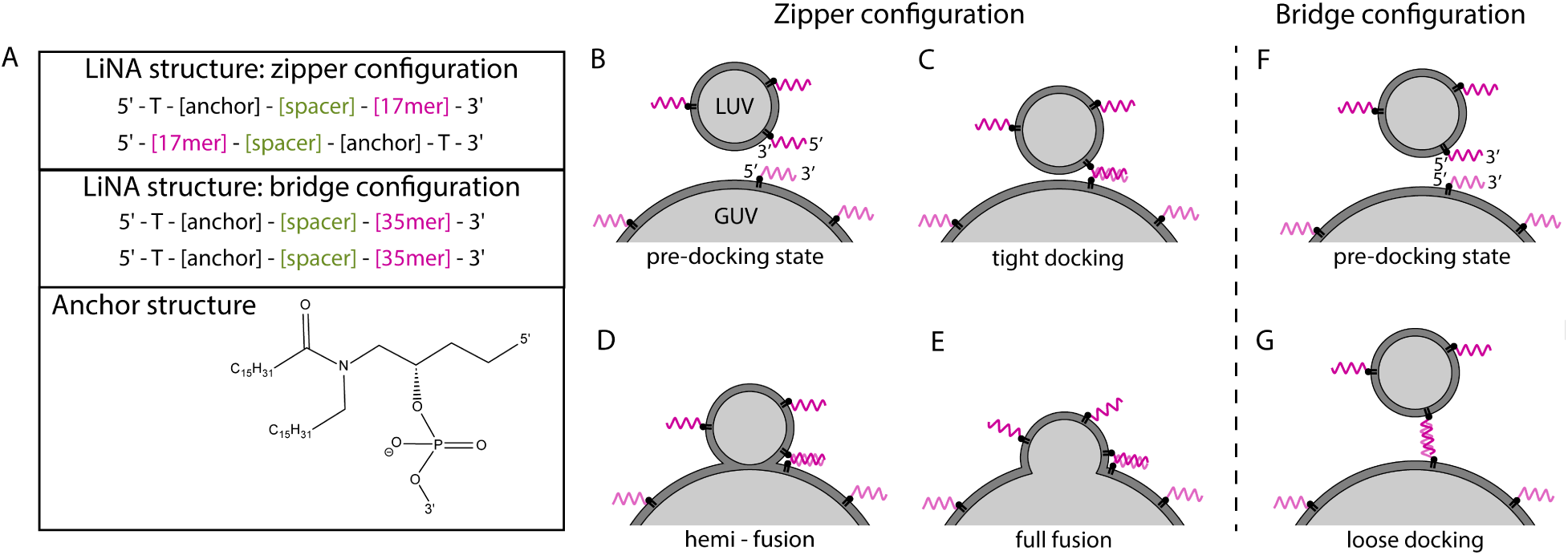
Schematic illustration of the programmable LiNA-mediated fusion approach. **A.** Overview of the molecular design of the hybridizing LiNA strands in the zipper configuration, which promotes LUV-to-GUV binding and fusion (top panel), and in the bridge configuration, which promotes binding but not fusion (middle panel). The configuration is determined by the placement of the membrane anchors on the oligonucleotides (bottom panel), on opposite ends in the zipper configuration and on the same end in the bridge configuration. LiNA sequences and the functionalization scheme for each vesicle type are provided in Supplementary Tables 1 and 2. **B-E.** Hybridization of complementary LiNA constructs in the zipper mode brings the GUV and LUV membranes in close contact (tight docking) and facilitates membrane remodeling events that - via a hemifusion intermediate state - result in full membrane fusion. **F-G.** LiNA constructs in the bridge configuration maintain an increased distance between the opposing GUV and LUV membranes upon hybridization, thereby resulting in loose docking but not in membrane fusion.

To validate that this zippering mechanism is essential to induce full membrane fusion, we employ a 35 nt LiNA construct with the anchors attached to the same ends of the DNA strands (both at the 5’ end), creating a bridge configuration (Figure 1F-G). Upon hybridization of these LiNAs, we anticipate that an increased distance (approximately 12 nm if the oligonucleotide is oriented perpendicularly to the membranes) is maintained between the opposing membranes that prevents membrane fusion. This situation was reported to tether vesicles together without inducing membrane fusion^28,31,58^ and therefore, we use this condition as a control condition to validate our lipid and content mixing assays.

### LiNAs efficiently mediate lipid mixing

The full merging of the GUV and LUV membranes is an essential prerequisite for full fusion. To assess membrane fusion, we employed a FRET assay based on fluorescence lifetime microscopy (FLIM), as this provides a sensitive readout of lipid exchange between the distinct vesicle populations (see schematic overview in Figure 2A.). We employ GUVs labeled with 0.1 mol% Atto 488-DOPE (donor) and LUVs labeled with 0.5 mol% Rho-DOPE (acceptor). Since the donor and acceptor fluorophores are confined to separate membranes and the FRET pair has a Förster radius (*R*_0_) of 5 - 6 nm^59^, no efficient energy transfer occurs when both vesicle populations coexist in solution. Docking of LUVs on the GUV membrane, probed by LiNAs in the bridge mode, should bring the opposing membranes into closer proximity and this may yield some energy transfer. Membrane fusion mediated by LiNAs in the zipper mode should merge the GUV and LUV membranes and enable efficient energy transfer between the donor and acceptor fluorophore as they are embedded within the same membrane. Here, FRET results in a reduction of the donor fluorescence lifetime as energy transfer from donor to acceptor accelerates donor decay, producing a shorter fluorescence lifetime. Indeed, monitoring the donor lifetimes (*τ*_d_) provides a robust, quantitative readout of membrane fusion that, importantly, is largely independent of absolute fluorophore concentration or intensity fluctuations. The use of cell-sized GUVs as a model system allows for direct detection of fusion at the single-GUV level by confocal FLIM imaging.

**Figure 2:**
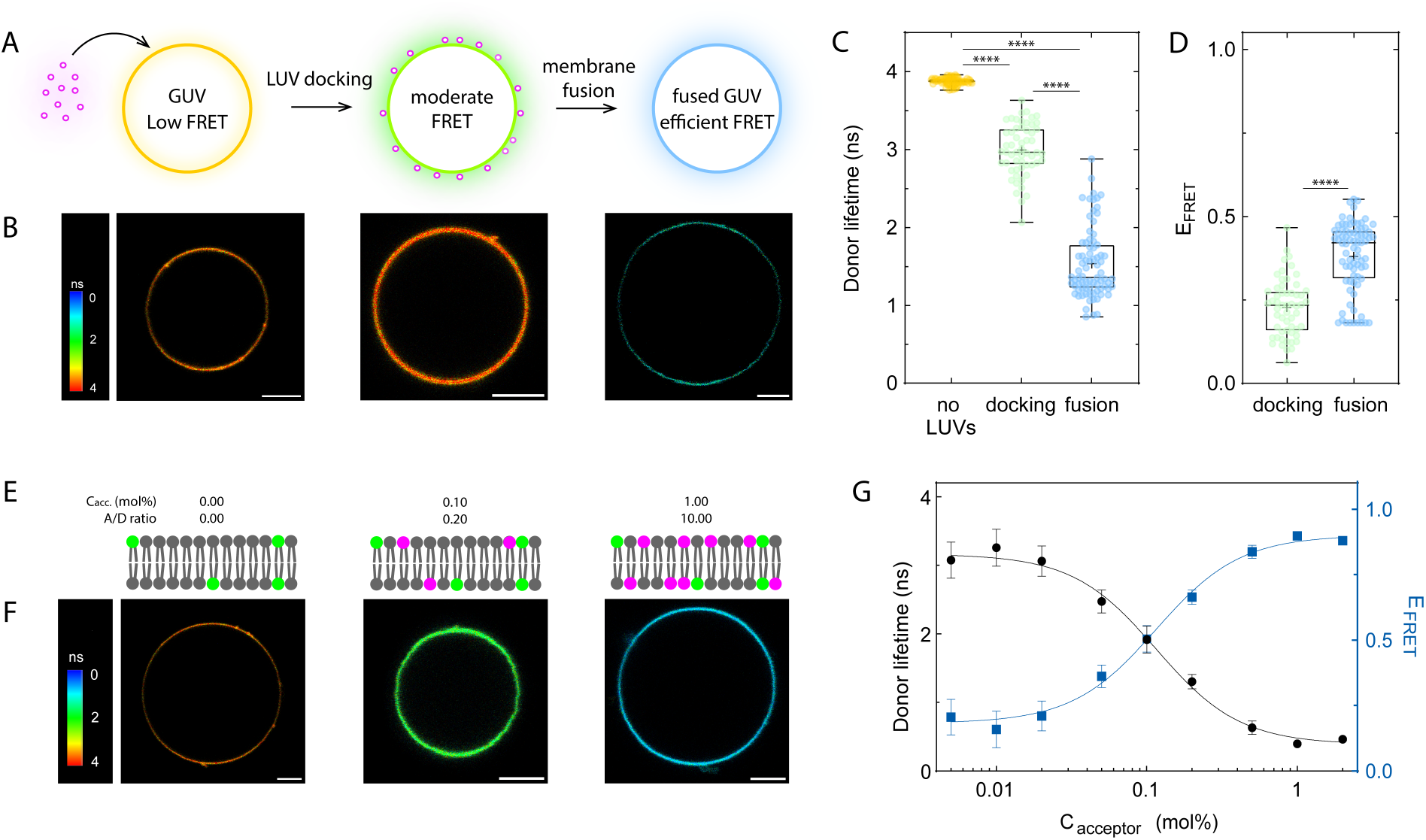
Lipid mixing assay to probe LUV-to-GUV fusion. **A.** Schematic representation of the lipid mixing assay. Fusion between GUVs labeled with donor dye (0.1 mol% Atto 488-DOPE) and LUVs labeled with acceptor dye (0.5 mol% Rho-DOPE) is monitored by measuring the change in fluorescence lifetime of the donor upon lipid mixing as a result of membrane fusion. **B.** Representative confocal FLIM images of GUVs in the absence of LUVs (left), after LUV docking mediated by 105 nM LiNA in the bridge configuration at 21°C (middle), and after fusion with LUVs mediated by 105 nM LiNA in the zipper configuration at 40 *^◦^*C (right). Confocal images are single slices taken at the GUV midplane. The color bar shows *τ*_d_. Scale bars are 5 *µ*m. **C.** Measured *τ*_d_ for GUVs in the absence of LUVs (left; from N = 3 independent repeats with n = 75 GUVs), after LUV docking mediated by LiNA in bridge mode (middle; N = 2, n = 56), and after fusion with LUVs mediated by LiNA in zipper mode (right; N = 3, n = 78). Statistical significance was tested using a non-parametric Kruskal-Wallis test (**** p *<* 0.0001). **D.** *E*_FRET_ values calculated from the *τ*_d_ measurements in panel C. Statistical significance was tested using a non-parametric Kruskal-Wallis test (**** p *<* 0.0001). **E.** Schematic representation of a series of GUV membranes containing different acceptor - donor (A/D) molar ratios to calibrate the dependence of *τ*_d_ on the concentration of acceptor dye in the GUV membrane. The donor dye is depicted in green, the acceptor dye in pink. **F.** Representative FLIM images of a series of GUVs containing different A/D ratios. Confocal images are single slices taken at the GUV midplane. The color bar shows *τ*_d_, scale bars are 5 *µ*m. **G.** Calibration curves showing measured donor fluorescence lifetime *τ*_d_ (black circles, left y-axis) and corresponding FRET efficiencies *E*_FRET_ (blue squares, right y-axis) as a function of the mole percentage of acceptor in the GUV membrane. Data are represented as mean ± SD.

First, we performed FLIM measurements on GUVs in the absence of LUVs. We measured a *τ*_d_ of 3.9 ± 0.05 ns (mean ± SD; image in Figure 2B first panel, quantification in Figure 2C). Representative decay curves are shown in Supplementary Figure 2. Then, we probed docking by functionalizing GUVs and LUVs with complementary LiNAs in the bridge conformation. After one hour of incubation at 21 *^◦^*C, we observed a *τ*_d_ of 3.0 ± 0.3 ns (Figure 2B middle panel, Figure 2C) and a corresponding FRET efficiency (*E*_FRET_) of 23 ± 9 % (Figure 2D).

The calculation of *E*_FRET_ is described in the Supplementary Methods. Subsequently, we probed fusion of LUVs with GUVs by complementary LiNAs in the zipper configuration. After one hour of incubation at 40 *^◦^*C, we measured a *τ*_d_ of 1.5 ± 0.5 ns (Figure 2B right panel, Figure 2C), corresponding to a *E*_FRET_ value of 38 ± 11 % after correcting for the contribution of docking (Figure 2D) as described in the supplementary methods. Lastly, we performed an additional control experiment in which we incubated unfunctionalized GUVs and LUVs for one hour at 40 *^◦^*C. We performed FLIM measurements and measured a *τ*_d_ of 3.8 ± 0.2 ns (Supplementary Figure 3).

Our observations show that tethering LUVs on the GUV membrane with LiNA in the bridge mode only induces docking, as is evidenced from a significant change in *τ*_d_ and in *E*_FRET_. This finding demonstrates that docking must be explicitly accounted for when calculating fusion efficiencies. Critically, when membrane fusion was probed using LiNA in the zipper configuration, a pronounced further reduction in *τ*_d_ and a concomitant increase in *E*_FRET_ were observed, consistent with the merging of the LUV and GUV membranes and the resulting co-localization of the donor-acceptor pair within a single bilayer. Taken together, these results provide robust fluorescence lifetime-based evidence for LiNA-mediated LUV-to-GUV fusion and highlight the importance of distinguishing docking from fusion when interpreting lipid mixing assay data.

To translate the observed *E*_FRET_ values into a fusion efficiency and calculate the number of LUVs that fused to a GUV, we performed a series of calibration measurements on GUVs containing a fixed amount of the FRET donor (0.1 mol%) and an increasing amount of the FRET acceptor (0.005 - 2 mol%) (Figure 2E, F). Representative decay curves are shown in Supplementary Figure 4. This enabled us to determine the relation between the measured *E*_FRET_ and the concentration of acceptor fluorophore in the GUV membrane (see Figure 2F). As LUVs have an average diameter of 160 nm (Supplementary Figure 5) and contain 0.5 mol% acceptor fluorophore, we determined that on average 280 LUVs fused to a single GUV with a radius of 10 *µ*m. By taking the GUV size into account (size distribution is shown in Supplementary Figure 6), we calculated a fusion density of 0.2 ± 0.06 events per *µm*^2^ of GUV membrane surface. The conversion of *E*_FRET_ into fusion density is described in the supplementary methods.

### Real-time observation of lipid mixing on single GUVs

To unravel the dynamics of the LiNA-mediated fusion process, we performed real-time measurements of lipid mixing on individual GUVs that were immobilized in a microfluidic trap (Figure 3A and Supplementary Figure 7). GUVs labeled with Atto 488-DOPE were first functionalized with 105 nM LiNAs and flushed through the device, where they encountered microposts that form a hydrodynamic trap. Once sufficient GUVs were stably immobilized by these traps, LUVs that were functionalized with complementary LiNA constructs and fluorescently labeled with Rho-DOPE were flushed through the device. A fresh batch of LUVs was added to the inlet every 15 minutes to prevent LUV depletion during the measurement. After 10 minutes, Rho-DOPE was detected on the surface of GUVs, indicating that LUVs were docking on the GUV membrane (Figure 3B). This was accompanied by a gradual reduction of *τ*_d_ over a time period of approximately 1 hour, after which it reached a plateau (Figure 3C). This observation indicates that membrane fusion mainly occurs within one hour. More specifically, we observed a *τ*_d_ of 3.5 ± 0.4 ns, 2.4 ± 0.7 ns, 1.8 ± 0.3 ns, 1.5 ± 0.4 ns, and 1.3 ± 0.4 ns at 15, 35, 65, 95, and 135 minutes after the start of LUV addition. These *τ*_d_ values correspond to *E*_FRET_ values (corrected for docking) of 0 %, 16 ± 19 %, 31 ± 9 %, 38 ± 11 %, and 45 ± 11 %, respectively (Figure 3C). We hence observe a delay between the first appearance of the LUV signal on GUVs and the increase of *E*_FRET_, which indicates that the transition from docking to lipid mixing took approximately 20 minutes.

**Figure 3:**
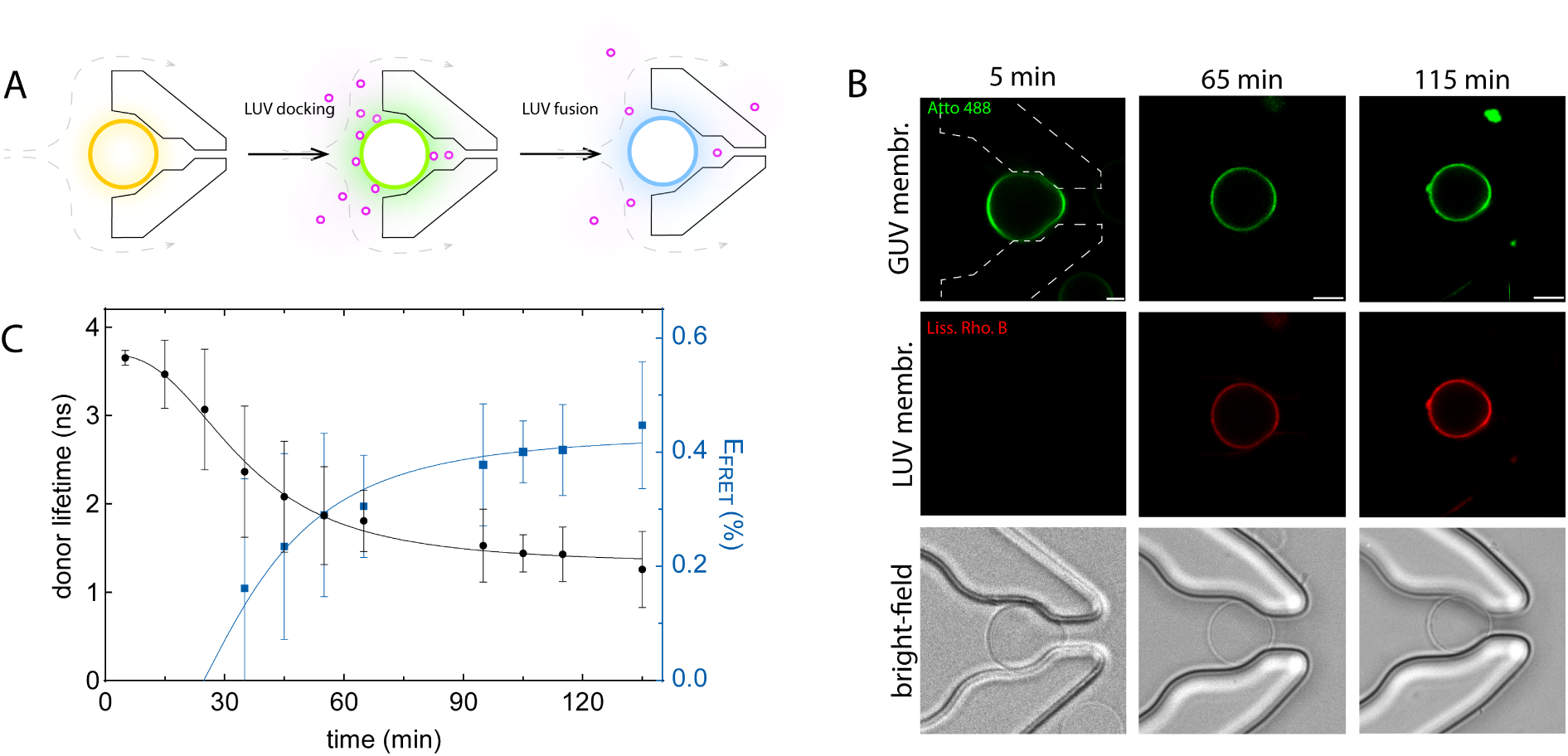
Real-time lipid mixing assay to probe GUV-LUV fusion using a microfluidic device. **A.** Schematic representation of the real-time lipid mixing assay. First, GUVs are drawn through the device and immobilized by microposts that form hydrodynamic traps (left). Then, LUVs are flushed through the device and dock on the GUV membrane upon hybridization of the LiNAs (middle). After docking, the fusion cascade proceeds further until membrane fusion occurs (right). The occurrence of fusion is monitored by measuring the fluorescence lifetime as in Figure 2A (see color code). **B.** Representative images of distinct GUVs at different time points after LUV addition. The GUV membrane is labeled with 0.1 mol% Atto 488-DOPE (green) and the LUV membrane is labeled with 0.5 mol% Rho-DOPE (red). The outline of a trapping structure is shown in the top-left panel in dashed white lines. Confocal images are single slices taken at the GUV midplane. The scale bar is 5 *µ*m. **B.** The evolution of *τ*_d_ (black circles, left y-axis) and *E*_FRET_ (grey squares, right y-axis) over time (N = 2, n = 29 for 5 min, n = 17 for 15 min, n = 23 for 25 min, n = 23 for 35 min, n = 23 for 45 min, n = 30 for 55 min, n = 17 for 65 min, n = 16 for 95 min, n = 7 for 105 min, n = 9 for 115 min, and n = 16 for 135 min). Data are shown as mean ± SD.

### LiNAs efficiently induce content mixing

To demonstrate that LiNA-mediated fusion results in full fusion, we quantified the mixing between the luminal contents of GUVs and LUVs. This is important and more compelling than the lipid mixing assay because membrane hemi-fusion, the intermediate state in which the outer but not the inner leaflets have merged, can result in lipid mixing and a measurable FRET signal in the previously described lipid mixing assay. A measurable FRET change could thus be erroneously taken as a false-positive indication of full fusion. We therefore set up a series of content mixing experiments in which non-fluorescent GUVs were fused to LUVs containing the dye sulforhodamine B (SRB) at a concentration of 20 mM (Figure 4A, left panel). When LUVs dock on the GUV surface, SRB is expected to be retained in the lumen of the LUVs and a SRB signal is only observed at the GUV surface (Figure 4A, middle panel). Full fusion is supposed to result in the mixing of the luminal contents of GUVs and LUVs, as evidenced by an increase of SRB fluorescence in the GUV lumen (Figure 4A, right panel). To correlate the measured SRB intensity to the SRB concentration (indicated as [SRB]) we performed a series of calibration measurements in which we measured the intensity of a concentration series of SRB solutions (supplementary Figure 8). The conversion of measured [SRB] in the GUV lumen after fusion into a calculated fusion density is described in the supplementary methods.

**Figure 4:**
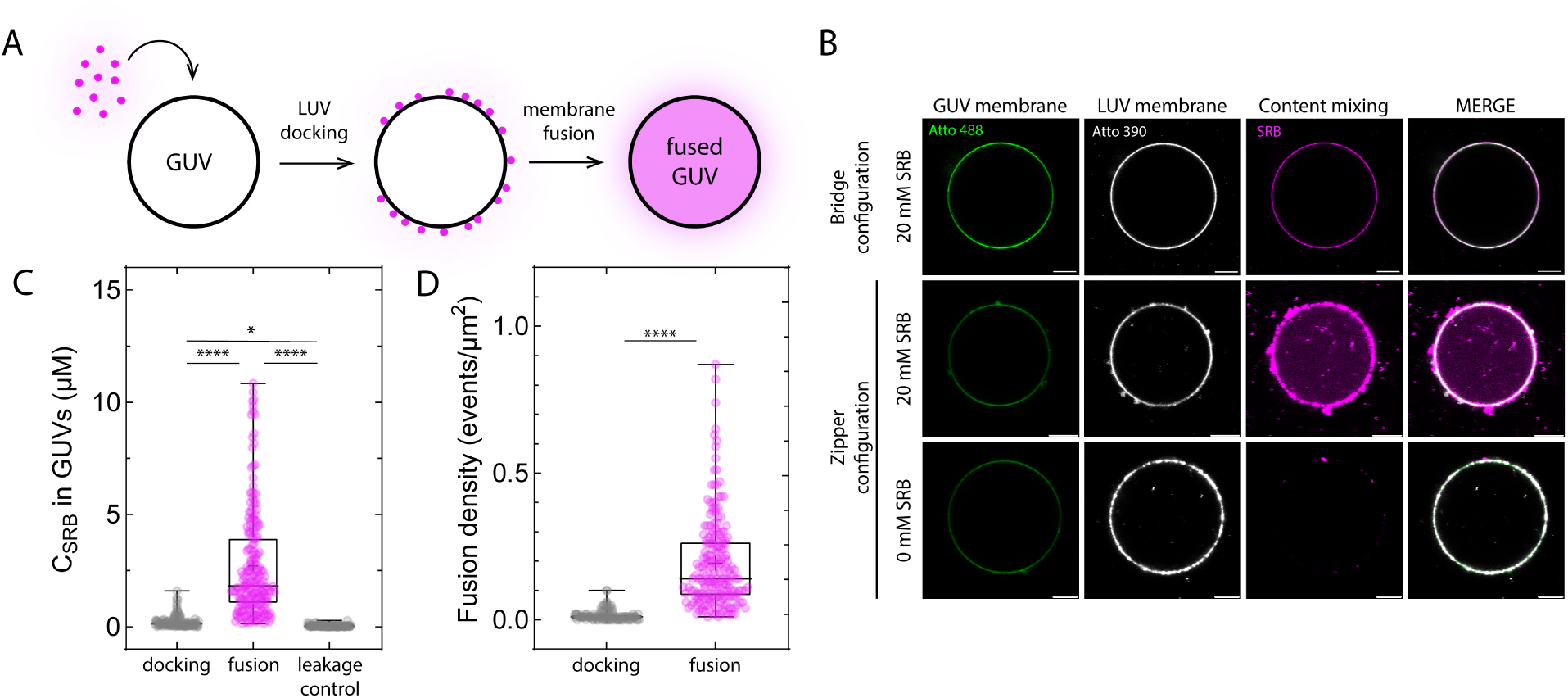
Content mixing assay to probe LUV-to-GUV fusion. **A.** Schematic representation of the content mixing assay. Fusion between empty GUVs and LUVs encapsulating 20 mM of the fluorescent dye SRB results in the transfer of the fluorescent dye into the GUV lumen, which can be observed by fluorescence microscopy. **B.** Representative confocal images depicting a GUV with LUVs docked to its surface mediated by LiNAs in the bridge configuration at 21 *^◦^*C (top row), a GUV that underwent fusion with LUVs containing 20 mM SRB mediated by LiNAs in the zipper configuration at 40 *^◦^*C (middle row), and a GUV that underwent fusion with empty LUVs mediated by LiNAs in the zipper configuration *^◦^*C (bottom row). The GUV membrane is labeled with Atto 488-DOPE (green), the LUV membrane is labeled with Atto 390-DOPE (white), and content mixing is observed by visualizing the transfer of SRB (magenta). Images are confocal slices at the GUV midplane. Scale-bars are 10 micron. **C.** Measured SRB concentration in the GUV lumen after the docking of LUVs encapsulating 20 mM SRB mediated by LiNAs in the bridge conformation (left; N = 4, n = 112) and after fusion with LUVs encapsulating 20 mM SRB mediated by LiNAs in the zipper conformation (middle;N = 6, n = 200) or with empty LUVs mediated by LiNAs in the zipper configuration (right; N = 4, n = 90). Statistical significance was tested using a non-parametric Kruskal-Wallis test (* p = 0.0239, **** p *<* 0.0001). **D.** Fusion densities based on the measured SRB concentrations shown in panel C. Statistical significance was tested using a non-parametric Kruskal-Wallis test (ns p *>* 0.9999, **** p *<* 0.0001).

When we functionalized GUVs and LUVs with 105 nM complementary LiNAs in the bridge configuration, we observed that LUVs localized on the GUV membrane. SRB, the marker for content mixing, also localized on the GUV surface, but no transfer into the GUV lumen was observed (Figure 4B top row). After one hour of incubation, however, we measured an apparent SRB concentration [SRB] of 0.2 ± 0.3 *µ*M in the GUV lumen (Figure 4C left), corresponding to a fusion density of 0.02 ± 0.02 events per *µm*^2^ of GUV membrane surface (Figure 4 D left). In contrast, functionalization of the vesicles with 105 nM complementary LiNAs in the zipper configuration resulted in the localization of the LUVs on the GUV surface as well as the transfer of SRB into the GUV lumen (Figure 4B, middle row). We measured an average [SRB] of 2.7 ± 2.4 *µ*M in the lumen of the GUVs (Figure 4C right), corresponding to a fusion density of 0.2 ± 0.2 events per *µm*^2^ of GUV membrane surface (Figure 4D right).

To verify that the observed SRB signal inside the GUV lumens originates from fusion and not from membrane destabilization and consequent dye leakage, we performed additional control measurements using empty LUVs functionalized with LiNA in zipper configuration. After formation of the LUVs, 20 mM free SRB dye was added to the external buffer solution. All subsequent sample handling steps were performed as described before. Confocal imaging showed that the LUVs localized on the GUV surface, but no SRB signal was detected in the GUV lumen (Figure 4B, bottom row). We measured a [SRB] of 0.04 ± 0.04 *µ*M in the GUV lumens (Figure 4C, right), which can be attributed to fluorescence background (Supplementary Figure 9).

These observations demonstrate that, when utilizing LiNAs in the bridge configuration, docking dominates over fusion and fusion events are rare. Conversely, when utilizing LiNAs in the zipper configuration, content mixing was clearly detectable, indicating that the contents of the LUVs and the GUV mix and thus providing evidence for full membrane fusion. This is in agreement with the results of our previously described FLIM-based lipid mixing assay. Importantly, our experiments can rule out a non-specific uptake of SRB due to leakage as a source of SRB signal in GUVs.

Taken together, we show that LiNA constructs in the zipper configuration efficiently mediate the mixing of the luminal contents of GUVs and LUVs and that this approach can be used for the delivery of small molecules on demand. We note that similar fusion densities were obtained from the lipid and content mixing assay, indicating that the vast majority of fusion events that are detected via both assays proceed to full membrane fusion.

### Determinants of LUV-to-GUV fusion

To obtain a better understanding of the mechanisms underlying LiNA-mediated membrane fusion, we explored several parameters that have been reported to influence membrane fusion: the incubation temperature, the oligonucleotide length of the LiNA construct, and the concentration of the LiNA constructs. All experiments described here are based on the lipid mixing assay described above. We used the *τ_d_* values as a metric for fusion to identify differences across conditions.

First, we tested the effect of the incubation temperature on lipid mixing. When GUVs and LUVs were functionalized with 105 nM complementary LiNAs in the bridge mode and incubated for one hour at 21 *^◦^*C, we measured a *τ_d_* of 3.0 ± 0.3 ns, whereas incubation at 40 *^◦^*C resulted in a *τ_d_* of 2.6 ± 0.5 ns (Figure 6A). Thus, LiNA constructs in the bridge mode do not mediate membrane fusion, even at elevated temperatures where thermal fluctuations enhance membrane contacts. When employing LiNAs in the zipper mode, however, we observed a strong decrease of *τ_d_* from 2.9 ± 0.4 ns when incubated at 21 *^◦^*C to 1.5 ± 0.5 ns when incubated at 40*^◦^*C (Figure 6A). Thus, LiNAs in the zipper mode promote lipid mixing in a temperature-dependent manner, consistent with previous reports showing that LiNA-mediated LUV-to-LUV fusion is strongly increased at elevated temperatures^37,38^. This means that LiNA-mediated fusion can be switched on or off at will by tuning the incubation temperature over a physiologically useful range between 20-40*^◦^*C.

Second, we tested the effect of oligonucleotide length, comparing the effect of a 17 nt and a 35 nt ssDNA oligonucleotide on lipid mixing. Using a temperature of 40 *^◦^*C and constructs in the zipper configuration, we observed comparable *τ_d_* values of 1.5 ± 0.5 ns for a 17mer LiNA and 1.5 ± 0.5 ns for a 35mer (Figure 5B). This indicates that the oligonucleotide length in this range does not have any appreciable effect on fusion efficiency. This observation is consistent with other reports and indicates that the release of free energy upon DNA hybridisation is not transmitted to the membrane^32^.

**Figure 5:**
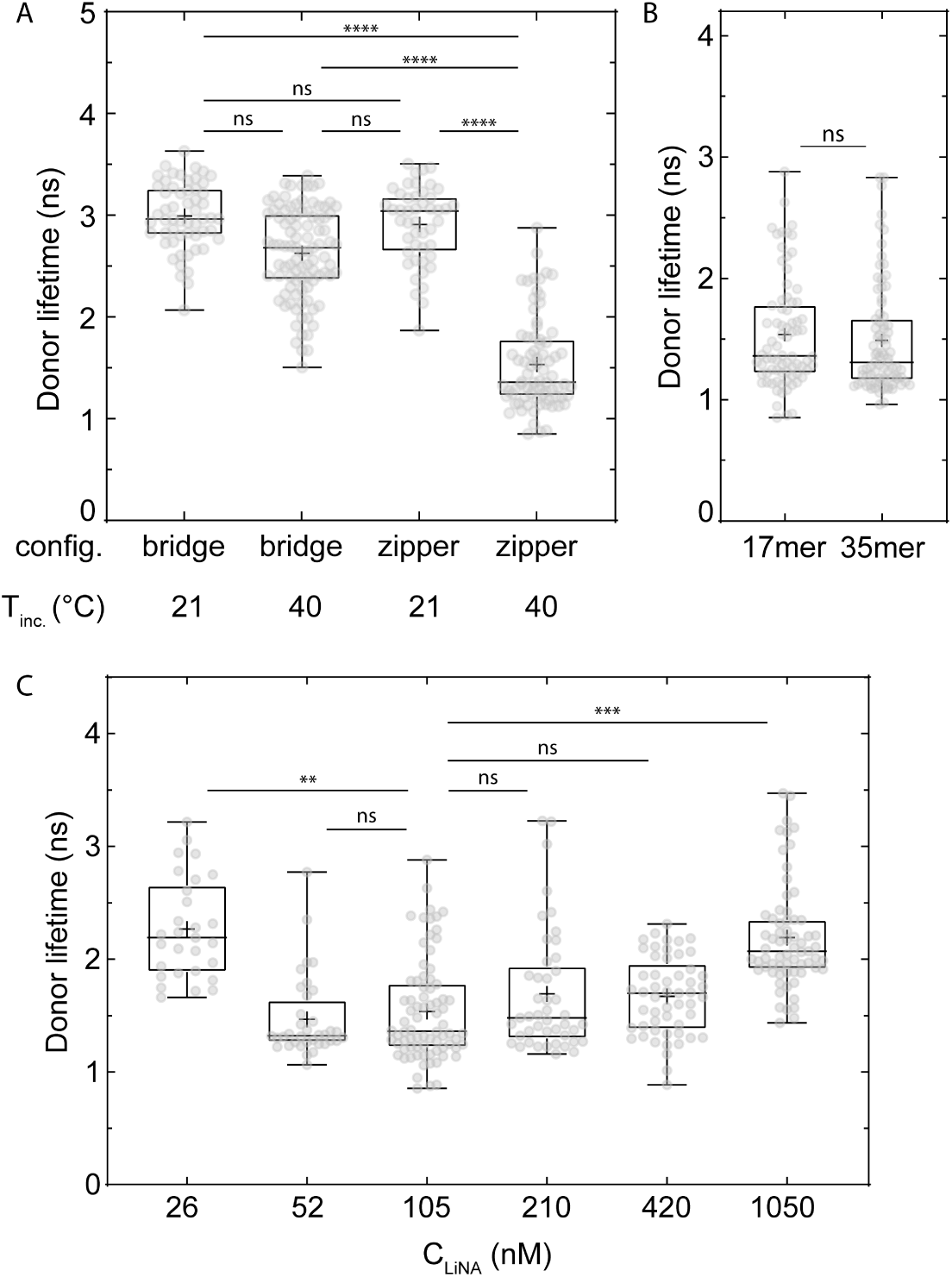
Determinants of LUV-to-GUV fusion. **A.** Effect of incubation temperature on lipid mixing. Measured *τ_d_* after incubation with 105 nM LiNA in the bridge mode at 21 °C (N = 2, n = 56) and 40 ◦C (N = 2, n = 96; left boxplots), and after incubation with 105 nM LiNA in the zipper mode at 21 ◦C (N = 2, n = 47) and 40 ◦C (N = 3, n = 78). Statistical significance was tested using a non-parametric Kruskal-Wallis test (**** P *<*0.0001, ns P *>* 0.9999). **B.** Effect of oligonucleotide length on lipid mixing. Measured *τ_d_* after incubation with 105 nM LiNA in the zipper mode at 40 *^◦^*C when employing a 17mer (N = 2, n = 82) and a 35mer (ns p *>* 0.9999). **C.** Effect of LiNA concentration on lipid mixing. Measured *τ_d_* after incubation with the 17mer LiNA in the zipper mode at 40 *^◦^*C. [LiNA] was 26 nM (N = 2, n = 29), 52 nM (N = 2, n = 36), 105 nM (N = 3, n = 78), 210 nM (N = 2, n = 45), 420 nM (N = 2, n = 53), 1050 nM (N = 2, n = 67). Statistical significance was tested using a non-parametric Kruskal-Wallis test (** P = 0.0099, *** P = 0.0004, ns P *>* 0.9999).

Third, we tested the impact of tuning the bulk concentration of the 17mer LiNA construct in the zipper configuration over a wide range of [LiNA], from 26 to 1050 nM, at a fixed temperature of 40 *^◦^*C (Figure6C). At the lowest LiNA concentration of 26 nM, we observed only a slight drop of *τ_d_* to 2.3 ± 0.4 ns, indicating a low degree of lipid mixing. In the [LiNA] range from 53 nM to 420 nM, we observed a markedly shorter *τ_d_*, but with little variation with [LiNA]. Specifically, we measured 1.5 ± 0.4 ns for 53 nM, 1.5 ± 0.5 ns for 105 nM, 1.7 ± 0.5 ns for 210 nM, and 1.7 ± 0.3 ns for 420 nM. Our observation of constant fusion efficiency above a threshold LiNA concentration is consistent with previous work showing that efficient membrane fusion requires multiple LiNAs at the contact site to cooperatively induce fusion^60^, similar to (SNARE) protein mediated fusion^61^. At the highest [LiNA] of 1050 nM, we observed an upturn of *τ_d_*, to 2.2 ± 0.5 ns, indicating a reduction in the degree of lipid mixing. In this case, a high GUV membrane coverage by LiNAs likely hinders fusion due to steric hindrance, which prevents direct membrane contact and thus fusion^42,60^. We conclude that an intermediate [LiNA] as used throughout this work, 105 nM, is a good choice since it is well above the threshold for robust membrane fusion while minimizing consumption of DNA.

### On-demand delivery of fascin triggers actin-network formation

The controlled and timed delivery of biomolecules into GUVs has thus far remained a open challenge in the synthetic cell field. Given the high efficiency and controllability of LiNA-mediated fusion, we wondered whether it is feasible to use this approach to deliver proteins to the lumen of GUVs. For this proof of concept, we turned to controlled bundling of actin filaments by delivery of fascin, a 56 kDa actin-binding protein that is known to efficiently bundle actin filaments^52^. This system provides a convenient experimental read-out, because fascin-actin bundles are highly rigid and therefore distinct from semiflexible actin filaments (Figure 6A). At the same time, this system is of direct interest for cell-free constitution of complex actin-based processes in synthetic cells such as active shape changes, motility, and division^62^. Fascin plays a pivotal role in organizing filamentous actin (F-actin) into tight, parallel bundles important for the formation of dynamic cellular protrusions such as filopodia and lamellipodia^63^. Its controlled introduction into a minimal cell-like compartment therefore represents a compelling model for studying cytoskeletal self-organization in a well-defined bottom-up system.

**Figure 6:**
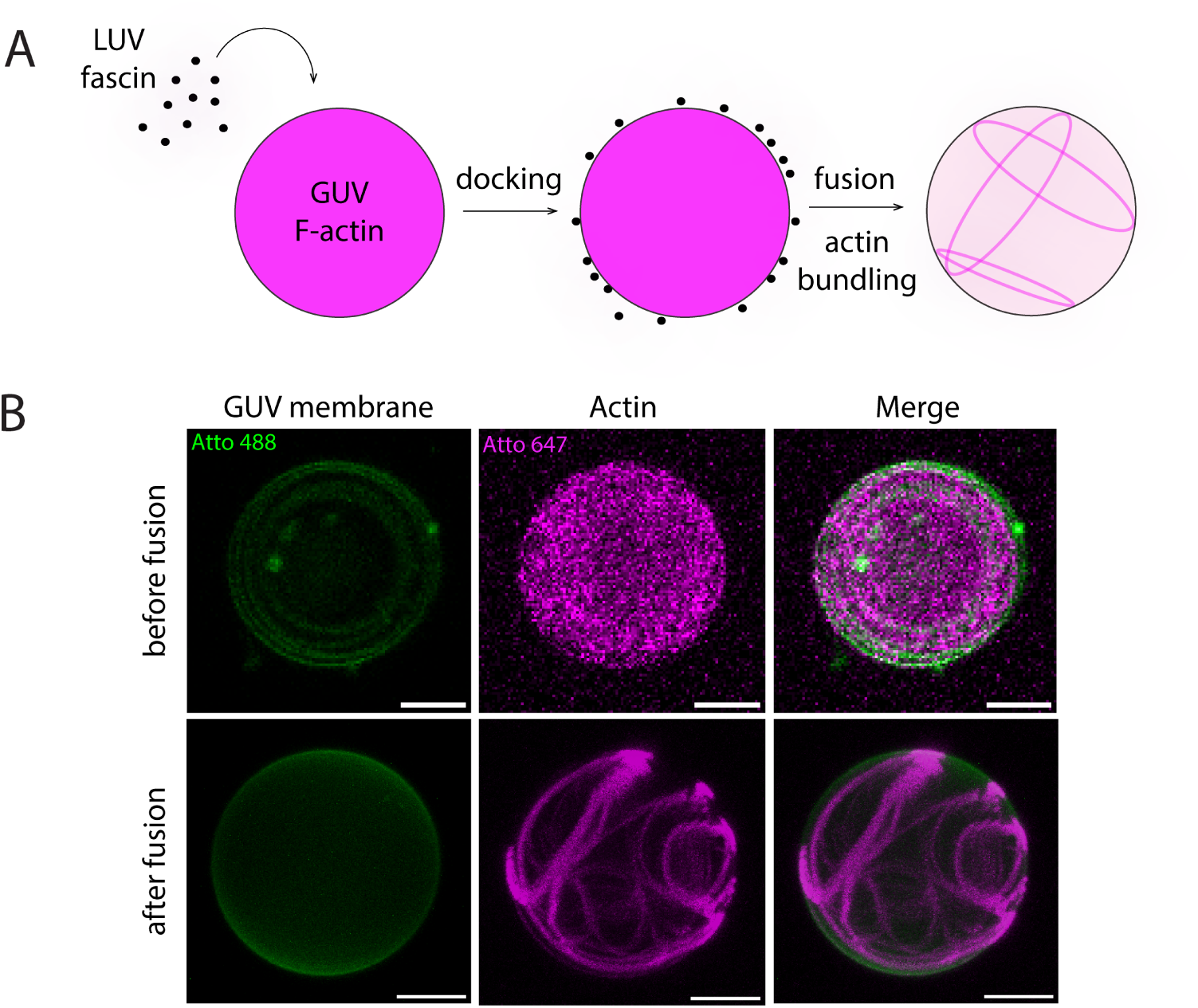
On-demand delivery of fascin by LUV-to-GUV fusion triggers bundling of actin filaments in the GUV lumen. **A.** Schematic representation of the assay to probe delivery of proteins into the GUV lumen. LUVs encapsulating fascin and GUVs encapsulating filamentous actin (F-actin; labeled with Atto 647) are functionalized with 105 nM complementary LiNAs in zipper configuration, mixed, and incubated for one hour at 40 °C. Upon fusion, fascin should be transferred into the GUV as detectable by a transformation from a homogeneous distribution of F-actin before fusion to a bundled F-actin network after fusion. **B.** Representative maximal intensity projections of GUVs encapsulating F-actin before (top row) and after fusion (bottom row) with LUVs. The delivery of fascin induces the formation of actin bundles. Columns show the GUV membrane labeled with Atto 488-DOPE (green; left), F-actin is labeled with Atto 647 (magenta; middle), and the merged images (right). Projections were recreated from z-stacks of 7 (top row) and 44 (bottom row) individual images spaced by 2.6 *µ*m (top row) and 0.6 *µ*m (bottom row), respectively. Scale bars are 5 *µ*m.

GUVs encapsulating prepolymerized F-actin and LUVs encapsulating fascin were each functionalized with 105 nM LiNA in the zipper configuration to promote membrane fusion. Prior to fusion, confocal imaging showed that F-actin was distributed homogeneously throughout the GUV lumen (Figure 6B, top row), consistent with the absence of bundling activity. After one hour of incubation at 40 *^◦^*C with the LUVs, however, we observed the formation of a bundled F-actin network, providing direct evidence that fascin was transferred into the GUV lumen. These networks were imaged by confocal microscopy, revealing the spatial organization of actin bundles within the GUV interior (Figure 6B). This proof-of-concept experiment underscores the power of LiNA-mediated LUV-to-GUV fusion as a versatile platform for on demand post-formation delivery of functional proteins to synthetic cells. Interestingly, the delivery can also be programmable and targeted to specific LUVs and/or GUVs, because the DNA sequence of the LiNA oligomers can be chosen at will and hence various LUVs with varying-sequence LiNAs can be chosen to fuse on demand^38,55^.

## Conclusions and outlook

In this work, we quantitatively demonstrated the successful establishment of on-demand LiNA-induced fusion between LUVs and micron-sized GUVs. To rigorously quantify fusion events that cannot be resolved by conventional optical microscopy, we developed two complementary assays to characterize lipid mixing and content mixing *in situ* at the single GUV level. This single-vesicle approach represents a significant methodological advancement over previous bulk measurements of LUV-to-LUV fusion, offering reduced interpretational ambiguity and greater experimental reliability. As our lipid and content mixing assays indicated a comparable extent of membrane fusion, we concluded that the fusion cascade primarily culminated in full membrane fusion, wherein both the membranes fused and the luminal contents mixed. Importantly, control experiments showed that this occurred without significant leakage. Further, we provided insights in the main factors governing membrane fusion. We demonstrated that the zipper conformation and elevated temperatures are essential for membrane fusion, whereas the DNA length of the constructs has no impact on fusion efficiency. As the GUVs employed in this study closely resemble cells in size and curvature, this system constitutes a minimal model system for studying membrane fusion *in vitro*.

Beyond providing mechanistic insight, we showed that the LiNA-mediated fusion system can be used for the targeted delivery of lipids as well as differently sized cytosolic components to synthetic cells. Thereby, this works extends the synthetic biologist’s toolbox to alter the composition of synthetic cells post-formation and trigger more complex reaction in a programmed manner. As a proof of concept, we delivered the actin-bundling protein fascin into GUVs containing F-actin, successfully triggering the formation of actin bundles.

The broader significance of the reconstitution and quantitative imaging assays described here extends well beyond the present study. Their versatility and robustness position them as powerful tools for a wide range of applications, including studying biomarker detection^64^, drug delivery^65^, or viral entry of the host cell^66^.

## Materials and Methods

### Materials

1,2-dioleoyl-sn-glycero-3-phosphocholine (DOPC), 1,2-dioleoyl-sn-glycero-3-phosphoethanolamine (DOPE), cholesterol (sheep wool), 1,2-dioleoyl-sn-glycero-3-phosphoethanolamine-N-(lissamine rhodamine B sulfonyl) (ammonium salt) (Rho-DOPE), and 1,2-dioleoyl-sn-glycero-3-phosphoethanolamine-N-[amino(polyethylene glycol)-2000] (ammonium salt) (PEG2K-DOPE) were obtained from Avanti Polar Lipids. DOPE labeled with Atto 390 and DOPE labeled with Atto 488 were obtained from ATTO-TEC. Lipid solutions were prepared in chloroform at a stock concentration of 25 mg/mL for non-headgroup-modified lipids, 10 mg/mL for PEG2K-DOPE, and 1 mg/mL for fluorescent lipids. Solutions were stored at -20 *^◦^*C until further use. Unlabeled *α*-actin and Atto 647-labeled *α*-actin (purified from rabbit skeletal muscle) were obtained from Hypermol. Recombinant fascin was purified in-house as described previously^52^. Chloroform, PVA, sucrose, glucose, NaCl, KCl, MgCl_2_, 2-[4-(2-hydroxyethyl)piperazine-1-yl]ethanesulfonic acid (HEPES), tris(hydroxymethyl)aminomethane-HCl (Tris-HCl), dithiothreitol (DTT), adenosine triphosphate (ATP), 3,4-protocatechuic acid (PCA), protocatechuate 3,4-dioxygenase (PCD), OptiPrep, SRB, Bovine serum albumine (BSA) were purchased from Sigma-Aldrich.

### Liposome Preparation

GUVs were prepared by the gel-assisted swelling method, since this method allows for complex lipid compositions including PE and cholesterol, needed for fusion^67^. Briefly, a 5%(w/v) (145 kDa, 98% hydrolysed) solution in MilliQ water was spread over a cover glass (24 x 24 mm; Menzel-Gläser) and baked at 50 *^◦^*C for 30 minutes to form a dried film. Then, 10 *µ*L of a lipid mixture (2 mM total lipid concentration; see Supplementary Table 2 for the exact lipid compositions) in chloroform was spread over the film and placed under vacuum for at least 30 minutes to evaporate the solvent. Subsequently, 300 *µ*L of swelling solution (100 mM sucrose, 100 mM NaCl, 10 mM HEPES at pH 7.0) was added on top of the film. The sample was then incubated at 21 *^◦^*C for 1 hour to allow GUV formation. Following incubation, the GUV solution was collected with a cut pipette tip to prevent disruption of the GUVs, stored at 4 *^◦^*C, and used within 48 hours of preparation.

For the encapsulation of proteins, we slightly adapted the gel-assisted method^67,68^. PVA was spin-coated on a round microscopy coverglass (30 mm diameter; Menzel-Gläser) at 4000 rpm for 30 s (Spin150, SPS Polos). 50 *µ*L of a lipid mixture (5 mM total lipid concentration; see Supplementary Table 2 for the exact lipid compositions) in chloroform was spin-coated on the dried PVA-coated cover glass at 1500 rpm for 300 s. A spacer with a thickness of 132 *µ*m (ARcare 90445Q, Adhesives Research) was placed on the PVA-coated side of the coverglass to form a chamber. Lipid swelling was initiated by the addition of 30 *µ*L swelling solution containing the proteins to be encapsulated (20 mM Tris-HCl (pH 7.4), 50 mM KCl, 2 mM MgCl_2_, 1 mM dithiothreitol (DTT), 1 mM adenosine triphosphate (ATP), 1 mM 3,4-protocatechuic acid (PCA), 0.05 *µ*M protocatechuate 3,4-dioxygenase (PCD), 6.5 %(v/v) OptiPrep, 5 *µ*M *α*-actin (10 % labeled)). PCA and PCD were used to reduce photobleaching, DTT was present to avoid actin oxidation, and ATP was present to promote actin polymerization. Optiprep was present to increase the density of the GUVs and gently sediment them to the bottom of the observation well. The chamber was then covered with another cover-glass and incubated for 45 min at 4 *^◦^*C to minimize actin polymerization during swelling. Then, 150 *µ*L of outer buffer (20 mM Tris-HCl (pH 7.4) supplemented with glucose to match the osmolarity of the inner solution, as verified using a freezing point osmometer(Gonotec Osmomat 3000)) was added to the chamber and GUVs were collected with a cut pipette tip. After harvesting the GUVs, actin polymerization was initiated by incubating the sample at 21 *^◦^*C. GUVs containing proteins were used on the day of preparation.

LUVs were prepared according to the thin film hydration method, followed by extrusion^69,70^. First, a lipid mixture (Supplementary Table 2) in chloroform was prepared in a 2 mL glass vial (Carl Roth). The solution was dried under vacuum for at least 30 minutes to ensure complete evaporation of the solvent. Then, the lipid film was hydrated by adding a swelling solution (100 mM glucose, 100 mM NaCl, 10 mM HEPES at pH 7.0) and vortexed to obtain a dispersion with a final total lipid concentration of 2 mM. This solution was then sonicated for 30 minutes in a bath sonicator (Branson 2510) at 21 *^◦^*C. To improve unilamellarity, the liposome solution was manually extruded 15 times (Avanti Mini Extruder, Avanti Polar Lipids) through a polycarbonate filter (Nucleopore, Whatman) with a pore size of 100 nm. The resulting LUV solutions were stored at 4 *^◦^*C and used within 48 hours of preparation. For content mixing experiments, the fluorescent dye sulforhodamine (SRB) was added to the swelling solution at a concentration of 20 mM. Unencapsulated dye was removed from the LUV solution by size exclusion chromatography (Zeba™ Dye and Biotin Removal Spin Column; ThermoFisher Scientific) following the manufacturer’s manual.

### Dynamic light scattering

The LUV size distribution was measured by dynamic light scattering (DLS) using a Zetasizer Nano ZS (Malvern Panalytical). Each sample, consisting of a 200 *µ*L of LUV solution with a nominal lipid concentration of 500 *µ*M, was measured three times at 25 *^◦^*C under an angle of 173 *^◦^* (backscatter) in a disposable cuvette (BRAND® 1.5 mL semi-micro cuvette). Each measurement consisted of at least 14 runs. The measured scattering intensity was converted into the intensity autocorrelation function and then into the particle size distribution using the Zetasizer Nano software (version 3.30).

### Liposome functionalization and fusion assays

To probe fusion, GUVs at a nominal lipid concentration of 46 *µ*M were incubated for 15 minutes at 21 *^◦^*C with the A or A II type LiNA (Supplementary Table 1 and 2). The LiNA concentration was 105 nM unless stated otherwise. Similarly, LUVs at a total lipid concentration of 0.138 *µ*M were incubated with the A’, A’rev, or A’rev II type LiNA (Supplementary Table 1 and 2). The LiNA concentration was 105 nM unless stated otherwise. Then, GUVs and LUVs were mixed in a 1:1 volumetric ratio and incubated at 21 *^◦^*C (to probe docking) or 40 *^◦^*C (to probe fusion) for 1 hour in a PCR block, after which the samples were transferred to a BSA-coated well with a No. 1.5 polymer coverslip bottom (18-well *µ*-slide, Ibidi). Each well was passivated by incubating for 30 minutes with a 5 mg/mL BSA-solution and then rinsed 3 times with MilliQ water and dried.

To probe docking of LUVs on the GUV surface, a similar protocol was followed. Importantly, in this case GUVs were functionalized with the A II type LiNA, whereas LUVs were functionalized with the A’rev II type LiNA (Supplementary Table 2), and incubation was performed at 21 *^◦^*C.

For control measurements in which one or both LiNA constructs were left out, their volume was replaced by buffer solution.

### Confocal microscopy and fluorescence lifetime imaging

To assess lipid mixing, we performed FLIM-FRET measurements on an inverted Leica Stellaris 8 FALCON laser scanning confocal microscope using LasX software (version 4.8.2). During imaging, a constant temperature was ensured by using a box incubator (Okolab). Samples were illuminated using a 63x glycerol immersion objective (HC PL APO CS2 63X/1.30, Leica). Atto 488 was excited at a wavelength of 497 nm using a white light laser (WLL) at a repetition rate of 20 MHz and 0.25 *µ*W as measured at the imaging plane. Brightness was optimized by adjusting the objective correction collar prior to each experiment. These settings minimized pile-up artifacts where more than 1 photon per pulse are detected. To ensure the acquisition of enough photons per pixel to obtain reliable decay fits without affecting the photon rate per pulse, we adjusted the number of line repetitions per image. The emission light was passed through a 103 *µ*m pinhole and collected in the spectral range between 507 nm and 570 nm using a HyDX1 detector operated in digital mode with an internal gain of 10. Rho-DOPE was excited at a wavelength of 573 nm using a WLL at 0.5 *µ*W. Emission light was collected in the between 583 nm and 700 nm using a HyDS2 detector operated in analog mode.

Content mixing was assessed by laser scanning confocal microscopy, performed on the same microscope equipped with the same objective. Atto 390 was excited at a wavelength of 405 nm using a solid-state diode (DMOD) laser at 20 *µ*W, Atto 488 was excited at a wavelength of 497 nm using a WLL at 1 *µ*W, and SRB was excited at a wavelength of 569 nm using a WLL at 1 *µ*W. For all excitation wavelengths, we used a scan speed of 400 Hz, resulting in a pixel dwell time of 2.8 *µ*s. The dyes were excited sequentially and emission light was passed through a 103 *µ*m pinhole before reaching the detectors. Atto 390 emmision light was collected in the spectral range of 414 - 480 nm by a HyD X1 detector operated in digital mode with an internal gain of 50, Atto 488 emission light was collected in the range of 507 - 560 nm by a HyD S2 detector in analog mode with an internal gain of 50, and SRB emission light was collected in the spectral range of 575 - 750 nm using a HyDS2 detector in counting mode with an internal gain of 70.

### Microfluidic Device Fabrication and operation

Microfluidic devices were fabricated by a combination of photo- and soft lithography, conducted in class 100 (ISO 5) cleanroom facilities. The surface of a four inch wafer was primed with hexamethyldisilazane (HMDS; ABCR GmbH) to enhance resist adhesion by spin coating (500 rpm for 5 s, 1000 rpm for 55 s) and baking at 200 *^◦^*C for 2 minutes. Next, the negative photoresist ARN 4400.05 (Allresist GmbH) was applied by spin coating (500 rpm for 5 s, 4000 rpm for 55 s) followed by a pre-bake at 90 *^◦^*C for 2 minutes. The *µ*MLA maskless laser writer (Heidelberg Instruments), equipped with a 365 nm laser, was used to write the desired patterns on the coated wafer with a dose of 16 *mJ/cm*^2^. After exposure, the wafer was post-baked at 100 *^◦^*C for 5 minutes. Next, the resist was developed for 85 s in MICROPOSIT MF-321 developer (Dow) with continuous agitation. Development was stopped by submerging the wafer in deionised water for 30 s while continuously being agitated.

The photoresist patterns were dry etched in the silicon wafer by a Bosch process with an inductive coupled plasma (ICP) deep reactive-ion etcher (Adixen AMS 100 I-speeder). The etching step was performed using *SF*_6_ at a flow rate of 200 standaard cubic centimeters per minute (sccm) for 7 s with the ICP power set to 2000 W and the capacitive coupled plasma (CCP) bias power disabled. During the passivation step *C*_4_*F*_8_ was used at a flow rate of 80 sccm for 2.5 s with the ICP power maintained at 2000 W. The CCP power was operated in chopped low frequency bias mode, alternating between 60 W for 10 ms and 0 W for 90 ms. During the process, the sample holder was located at 200 mm from the plasma source, the He backside flow pressure was 10 mbar, the temperature of the wafer was kept at 10 *^◦^*C, and the temperature of the main chamber was 200 *^◦^*C. The photoresist was then stripped away in the same machine with the ICP power set at 2000 W with a biased power of 30 W, a source-target distance of 200 mm, temperature set at 10 *^◦^*C, using an *O*_2_ gas flow at 200 sccm for 5 min.

Structure heights were measured using a DektakXT (Bruker) stylus profiler equiped with a stylus with a radius of 0.7*µ*m. The wafer was then rendered hydrophobic by exposing it to (tridecafluoro-1,1,2,2-tetrahydrooctyl) trichlorosilane (ABCR GmbH) in a vacuum desiccator for at least 30 minutes.

PDMS microfluidic devices were fabricated by mixing the Sylgard 184 silicone elastomer base and curing agent (DOW) in a 10:1 weight ratio in a planetary mixer (Thinky ARE 250). The PDMS mixture was then poured over the wafer, degassed in a vacuum desiccator until no bubbles were visible anymore, and baked at 85 *^◦^*C for 4 hours. Then, the PDMS slab was peeled from the wafer and the devices were cut in the appropriate shape. Inlets (3 mm ID) and outlets (0.5 mm ID) were punched with a biopsy punch (World Precision Instruments). Microscopy slides (No. 1) and the PDMS blocks were placed in the chamber of a plasma reactor (Atto, Diener electronic GmbH + Co. KG) and exposed to an oxygen plasma at 50 W for 20 s to activate the surfaces. Immediately afterwards, the PDMS slabs were placed on the slides and baked on a hotplate at 130 *^◦^*C for 5 minutes to improve bonding.

Microfluidic devices were connected via Tygon Microbore tubing (0.020’ ID x 0.060’ OD; Saint-Gobain ICS) to a 250 *µ*L glass syringe (SETonic GmbH) that was operated with a Nemesys S syringepump (Cetoni GmbH) using the Cetoni Elements software (version 20230130).

First, the device was flushed with experimental buffer to remove air. Then, 10 *µ*L of GUV solution was added to the inlet, and flown through the device at a rate of 5 *µ*L/h until sufficient GUVs were collected in the traps. Then, 10 *µ*L LUV solution was added to the inlet, after which imaging was started immediately. During image acquisition, the flow rate was kept constant at 5 *µ*L/h. Every 15 minutes 10 *µ*L LUV solution was added again to the inlet to prevent LUV depletion.

### Data analysis and statistics

Fluorescence lifetime analysis was performed using the single molecule wizard of the Leica LasX software. The membrane signal at the equatorial plane of the GUV was manually selected. Photon arrival times were fitted with an n-exponential reconvolution function with n = 1 for donor only samples and n = 2 for all other samples. The instrument response function (IRF) generated by the software was used. The amplitude-weighted lifetimes were used for calculating *E*_FRET_. Conversion of the *E*_FRET_ in fusion densities is described in the Supplementary Methods.

Measurements of the SRB signal in the GUV lumen ass well as the GUV radius were performed using Fiji^71^ and a custom written Python script. Intensity thresholding was first applied to images of isolated GUVs, yielding a ring-shaped binary mask that delineated the fluorescent GUV membrane. This ring mask was cleaned and filled to produce a solid mask isolating the GUV area from the surrounding background. An inner mask, excluding the membrane signal and encompassing only the lumen, was then obtained by erosion of the solid mask. The mean fluorescence intensity within this inner mask was taken as the SRB lumen signal. Next, the inner mask was subtracted from the solid mask to yield a smooth ring mask isolating the membrane signal. Radial sampling of this masked edge image about its center of mass produced an [angle, radius] polar map, in which the membrane appeared as an approximately horizontal peak intensity line. The mean radial position of this peak, averaged across all angles, was taken as the GUV radius.

Values for *τ*_d_, *E*_FRET_, [SRB], and fusion densities were plotted in Graphpad Prism (version 10.6.1). All statistical analysis were performed in the same software. Prior to statistical comparison, the data were evaluated for normality and homogeneity of variance (Shapiro-Wilk test). As these conditions were not met, we used a non-parametric Kruskal–Wallis ANOVA test followed by a Dunn’s multiple comparisons test to evaluate the statistical significance between the samples.

## Supporting Information Available

Additional experimental methods, calculations, supplementary figures (Figure S1-11), and supplementary tables (Table S1 and 2) (PDF).

## Acknowledgement

We gratefully acknowledge financial support from The Netherlands Organization of Scientific Research (NWO/OCW) Gravitation Program Building a Synthetic Cell (BaSyC) (024.003.019). We furthermore thank Marcos Arribas Perez, Charu Sharma, Nikki Nafar, and Rafael B. Lira for advise and discussions regarding the experiments and data analysis. We thank Jeffrey den Haan and Sonam Marapin for protein purification and the Kavli Nanolab Delft support staff for their guidance on the fabrication of the microfluidic device.

## Author Contributions

Conceptualization: B.V.H, C.D., G.H.K.; investigation and formal analysis: B.v.H; Code development: J.K.; LiNA synthesis: N.A.R., S.V.; writing - original draft: B.v.H; writing - review and editing: B.v.H, S.V., C.D., G.H.K; funding acquisition: C.D., G.H.K.

## Data availability

The raw data and processed data for reproducing the figures are deposited in Zenodo (10.5281/zenodo.19316082) and will be made publicly available upon publication of the manuscript.

