## Supporting Information for "Lipid-conjugated DNA enables on-demand delivery of lipids and proteins to synthetic cells"

### Supporting Information Available

### Supplementary data and methods

#### Design, synthesis and characterization of the lipidated oligonucleotides

Two sets of LiNA were designed with different recognition lengths, 17 bp and 35 bp respectively. For both sets, a complementary pair (A+A' or AII+A'II) promotes fusion while the reversed strand hybridizes to form and locked bridge corresponding to the LiNA length (A+A'rev or AII+A'revII)[1]. The sequence design including structure of modifications is described in previous studies[2,3], and the XN lipid modification was synthesized according to the published method[4].

The LiNAs were synthesized using standard conditions for solid-phase synthesis. XN and P3 modifications were introduced as phosphoramidites. The P3 spacer was used as a 0.1 M solution in acetonitrile, and the XN modification was used as a 50 mM solution in a mixture of 1,2-dichlorethane and acetonitrile (2:1, v/v). Both modifications were hand-coupled to the sequence using a syringe with 0.4 mL of the amidite and 0.6 mL of Activator 42. The mixture was flushed through the column twice over a total time of 15 min for P3 and 25 min for XN. After synthesis, the oligonucleotides were cleaved from the solid support using ammonia at 55 °C for 24 hours. The solution was filtered, the ammonia was evaporated, and the oligonucleotide was re-dissolved in 50% acetonitrile and water (v/v). Purification was performed using reverse-phase HPLC (Waters, XBridge BEH C8, 5 µm, 130 Å), with solvent A = 0.05 M TEAA, solvent B = 0.05 M TEAA/acetonitrile (1:3 v/v). The HPLC method used a gradient of 10%-100% B followed by continuous flow of 100% B for at least 10 minutes to ensure elution of the lipidated oligos. All fractions were tested with matrix-assisted laser desorption/ionization (MALDI) mass spectrometry to verify the purity. Fractions containing the product were pooled and final tested for purity with analytical HPLC and MALDI followed by evaporation to dryness. All LiNA sequences and the MALDI results are summarized in Supplemntary Table 1.

LiNA stock solutions (10 $\mu$M) were stored in glass vials in a 50% acetonitrile and water (v/v) at 4 °C. At the start of each experiment, stock solutions were diluted in the experimental buffer to obtain a final concentration of 1 $\mu$M. These dilutions were stored at 4 °C and used up to 24 hours.

### Tm measurements of zipper and bridge LiNAs

The Tm for both 17mer and 35mer LiNAs are tested using UV measurements by a Cary 100 Bio Spectrophotometer. 250 nM of each LiNA was incubated with 250 $\mu$M POPC LUVs for 15 minutes at 37 °C. Corresponding LiNA pairs were then mixed in cuvettes and the UV absorbance was measured over 8 temperature ramps from 20-70 °C (A+A' and A+A'rev) and 20-80 °C (AII+A'II and AII+A'revII). Data are shown is Supplementary Figure10. For all experiments, the 1 ramp is an anealling step with 10 °C/min heating followed by 0.5 °C/min in the following ramps.

### SRB fusion assay for 100 nm LUV liposomes

An SRB fusion assay was performed for A+A' and A+A'rev to confirm that the bridge design does not promote fusion or content mixing. Two sets of 100 nm LUVs containing 20 mM SRB in one batch and PBS buffer in the second batch were extruded from a fusogenic lipid mixture of 28.5:45:25:1:0.5 (DOPC:DOPE:Chol:DOPG:DOPE-PEG2000). For LiNA A 200 nM LiNA is added to a 100 $\mu$M concentration of the SRB loaded fusogenic liposomes in a total volume of 200 $\mu$L. For complementary LiNAs A' and A'rev 200 nM LiNA is added to 100 $\mu$M of the empty PBS liposomes. The LiNA and liposomes are incubated for 20 minutes at 37 °C before all SRB loaded liposomes are transferred to cuvettes and the fluorescence measurements are started using a Cary Eclipse. Upon timepoint 0, the second population of PBS liposomes is added to the cuvettes, and the relative increase (I/I0) over time is plotted. For leakage experiments, two populations of SRB loaded liposomes are incubated with a complementary A+A' LiNA pair leading to no dilution upon fusion. The same control is performed for leakage upon assembly with A+A'rev. Data are shown is Supplementary Figure11 The results demonstrate content mixing for the zipper design of A+A' while the bridge design have a similar increase compared to the passive leakage controls.

### FLIM and calculation of FRET efficiencies

We used the organic dye Atto 488 as a FRET donor that displays a mono exponential decay. When energy transfer occurs, we observe a multi-exponential decay. Thus, donor decay functions are approximated by a bi-exponential model:

$$I(t) = I_0(A_1 exp(-t/\tau_1) + A_2 exp(-t/\tau_2)), \tag{1}$$

with $I(t)$ the intensity at time t, $I_0$ the intensity at time t=0, and $A_1$ and $A_2$ pre-exponential factors associated with $\tau_1$ and $\tau_2$. By using this model, amplitude-weighted lifetimes are given by:

$$\langle \tau \rangle_{\text{amp,DA}} = a_1\tau_1 + a_2\tau_2, \tag{2}$$

where $a_1$ and $a_2$ are the amplitudes of the lifetime components $\tau_1$ and $\tau_2$ and $a_1 + a_2 = 1$. The resulting $E_{\text{FRET}}$ was then calculated as follows:

$$E = 1 - \frac{\langle \tau \rangle_{\text{amp,DA}}}{\tau_D} \tag{3}$$

However, it is important to point out that $E_{\text{FRET}}$ as described in equation 3 overestimates the degree of membrane fusion. Both docking and membrane fusion result in FRET as shown in Figure 2C, D. Therefore, the measured FRET efficiency ($E_{\text{measured}}$) is a combination of FRET due to membrane fusion ($E_{\text{fusion}}$) and due to docking ($E_{\text{docking}}$):

$$E_{\text{measured}} = E_{\text{fusion}} + E_{\text{docking}} \tag{4}$$

To account for this and only consider FRET resulting from membrane fusion, we define the corrected FRET efficiency ($E_{\text{corrected}}$) that takes the contribution of docking to the total FRET signal into account:

$$E_{\text{corrected}} = E_{\text{measured}} - E_{\text{docking}} \tag{5}$$

For simplicity, ($E_{\text{corrected}}$) is always considered in the main text and used for calculating the extent of membrane fusion.

### Calibration of the FRET signal

To determine the number of fusion events based on the lipid mixing assay, $E_{\text{corrected}}$ determined as described above is converted to the mole fraction of the acceptor fluorophore ($\chi_{acc}$) present in the GUV membrane. Therefore, we performed a series of calibration measurements in which we used GUVs with a fixed mole fraction of the donor fluorophore ($\chi_{donor}$0.1 mol%) and an increasing $\chi_{acc}$ (0.005 - 2 mol% Rho-PE). As expected, the relationship between $E_{\text{FRET}}$ and $\chi_{acc}$ clearly showed a sigmoidal behavior (Figure 2G). We performed a sigmoidal 4 parameter logistic fit using the GraphPad Prism software (version11.0.0) which yielded the following relationship:

$$\chi_{acc} = 0.1155(\frac{E - 0.1824}{0.9000 - E})^{1/1.554}, \tag{6}$$

with $E$ taken as $E_{\text{corrected}}$ and a $R^2$ value of 0.9731. This relationship allows us to determine $\chi_{acc}$ in the GUV membrane after fusion on the single GUV level.

### Determination of fusion density based on lipid mixing

To determine the number of LUVs that fused to a single GUV, we measured $\tau_d$ in the GUV membrane (Figure 2B, C), converted this to $E_{\text{FRET}}$ (Figure 2D, Supplementary methods), and quantified $\chi_{acc}$ in the GUV membrane (Figure 2G, Supplementary methods). Additionally, the confocal images are single slices acquired at the equatorial plane of each GUV, allowing for the estimation of the average total GUV surface area (assuming each GUV to be a perfect sphere) as well as the total average number of lipids in the GUV membrane (assuming that each lipid occupies 0.65 nm[25,6]). Subsequently we determined $\chi_{acc}$ in the GUV membrane after fusion. Since the initial lipid mixture that was used for the preparation of the GUVs did not contain the acceptor dye, the only source of the acceptor dye in the GUV membrane are LUVs that have fused to the GUV.

We determined the average LUV surface area (80424.8 nm$^2$) from DLS measurements (Supplementary Figure 4). Using an estimated lipid head group area of 0.65 nm[25,6], we calculated that each LUV bilayer contains approximately 247461 lipids. Given that $\chi_{acc}$ in the membrane is 0.5 mol%, we determined that each LUV contains an average of 1237 acceptor dye molecules.

After having determined $\chi_{acc}$ in the LUV membrane before fusion and $\chi_{acc}$ in the GUV membrane after fusion, we are able to quantify the amount of fusion events that occurred for a single GUV. Normalization of the number of fusion events by the GUV surface area yields the average fusion density (fusion events/$\mu m^2$ GUV membrane surface).

#### Determination of fusion density based on content mixing

Similar to the lipid mixing assay, we also set up a methodology for calculating the fusion density based on our content mixing assay. We started by acquiring confocal images (single slices acquired at the equatorial plane) of each GUV after fusion. This allows for determination of the total GUV volume (again assuming that each GUV is a perfect sphere) and the SRB intensity in the GUV lumens. As the GUV initially do not contain any SRB and we can rule out leakage, the only source of SRB is from fusion with LUVs. Then, we converted the measured SRB intensity into the SRB concentration (Figure 4C, Supplementary Figure8) and the amount of SRB moles.

Each LUV encapsulates 20 mM SRB, has an average volume of 2.1 E$^{-18}L$, and contains a known amount of moles of SRB (4.29 E$^{-20}$). As each fusion event contributes a fixed amount of SRB to the GUV lumen, the amount of moles of SRB in the GUV lumen reflects the number of LUVs that have fused to it. By taking the GUV size into account we determined the fusion density (Figure 4D).

### Supplementary figures and tables

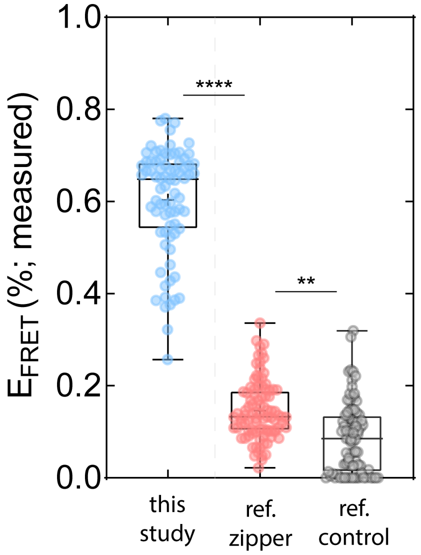


Figure 1: **Effect of membrane anchor structure on lipid mixing.** We assessed membrane fusion via lipid mixing and we observed significant differences in fusion efficiency when employing the LiNA constructs in zipper mode that are presented in this study and commercially available cholesterol-conjugated DNA constructs in zipper mode (constructs and experimental conditions were taken from Paez-Perez et al.[7]). For the LiNA constructs in zipper mode presented here, we measured a $E_{\mathrm{FRET}}$ of 60.3 ± 11.6 % (mean ± SD; from N = 3 independent repeats with n = 78 GUVs) after incubation for one hour at 40 °C, whereas a $E_{\mathrm{FRET}}$ of 14.7 ± 6.4 % (N = 3, n = 93) was measured for the reference construct in zipper mode after incubation for one hour at 21 °C. Furthermore, a $E_{\mathrm{FRET}}$ of 8.6 ± 7.8 % (N = 3, n = 84) was measured in a control experiment in which GUVs and LUVs were mixed without prior functionalization with DNA constructs, indicating that the DNA constructs only induce a minor degree of FRET and thus fusion. Statistical significance was tested using a non-parametric Kruskal-Wallis test (**** $p < 0.0001$, ** $p = 0.0013$).

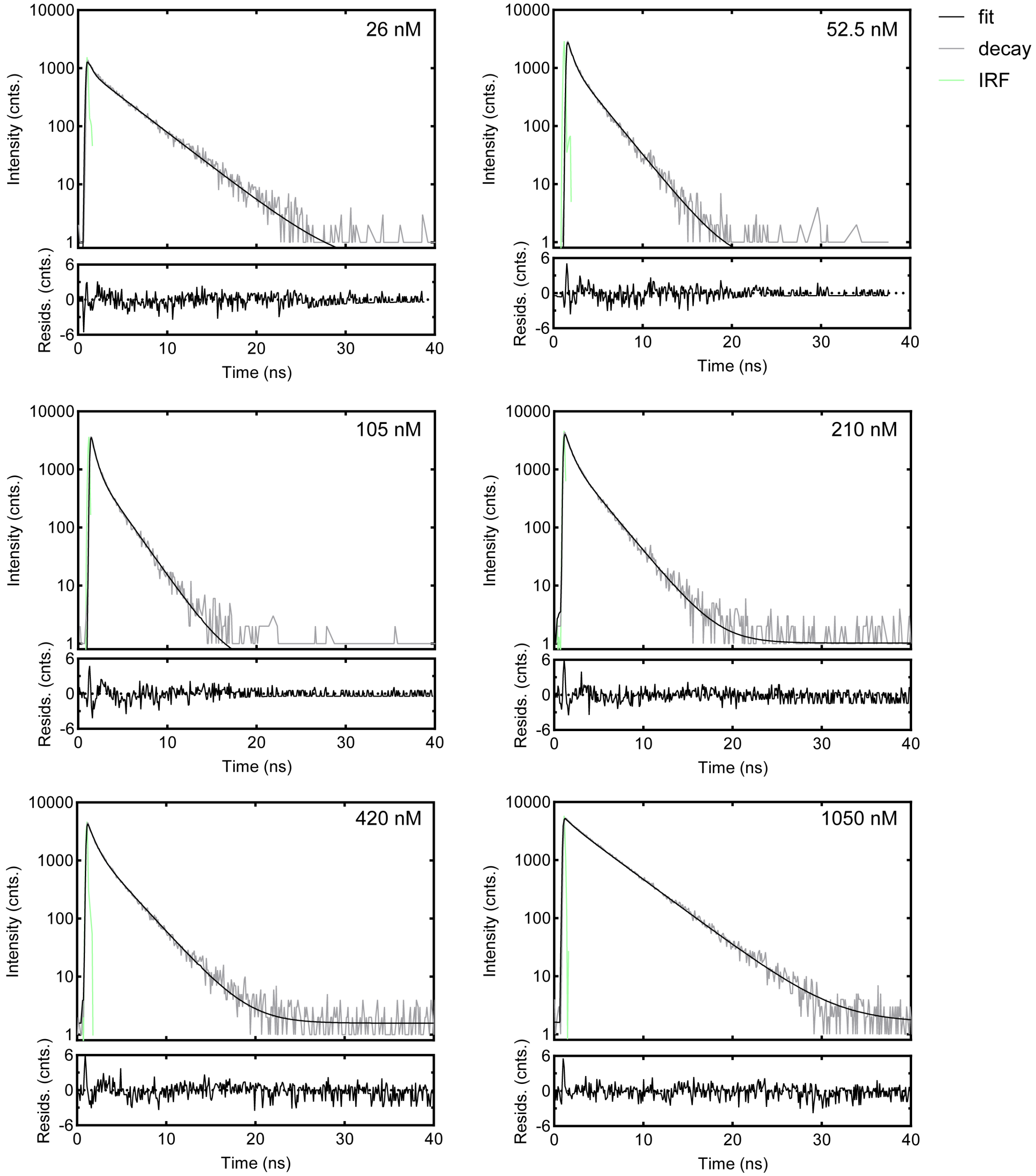


Figure 2: Representative FLIM decay curves for GUVs that fused with LUVs in the presence of different zipper LiNA concentrations (26 nM, 52.5 nM, 105 nM, 210 nM, 420 nM, 1050 nM) after one hour incubation at 40 °C. The decay curve is shown in gray, the fit is shown in black, and the instrument response function (IRF) is shown in green. The concomitant residual counts are plotted below each curve.

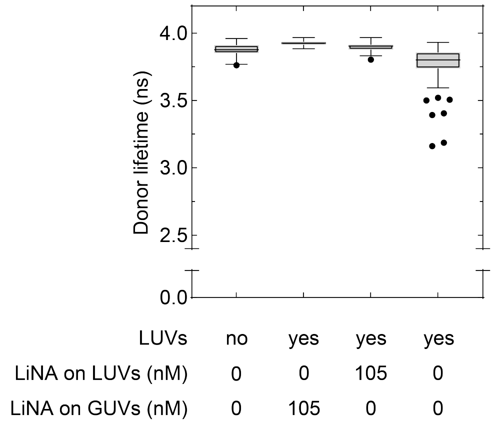


Figure 3: FLIM data for different control samples, showing measured $\tau_d$ values. Unfunctionalized GUVs in the absence of LUVs had a $\tau_d$ of 3.9 ± 0.05 ns (N = 3, n = 75). Functionalized GUVs in the presence of unfunctionalized LUVs had a $\tau_d$ of 3.9 ± 0.02 ns (N = 2, n = 20). Unfunctionalized GUVs in the presence of functionalized LUVs had a $\tau_d$ of 3.9 ± 0.04 ns (N = 2, n = 20). Unfunctionalized GUVs in the presence of unfunctionalized LUVs had a $\tau_d$ of 3.8 ± 0.2 ns (N = 3, n = 61). ns from P > 0.9999 in a Dunn's multiple comparisons test.

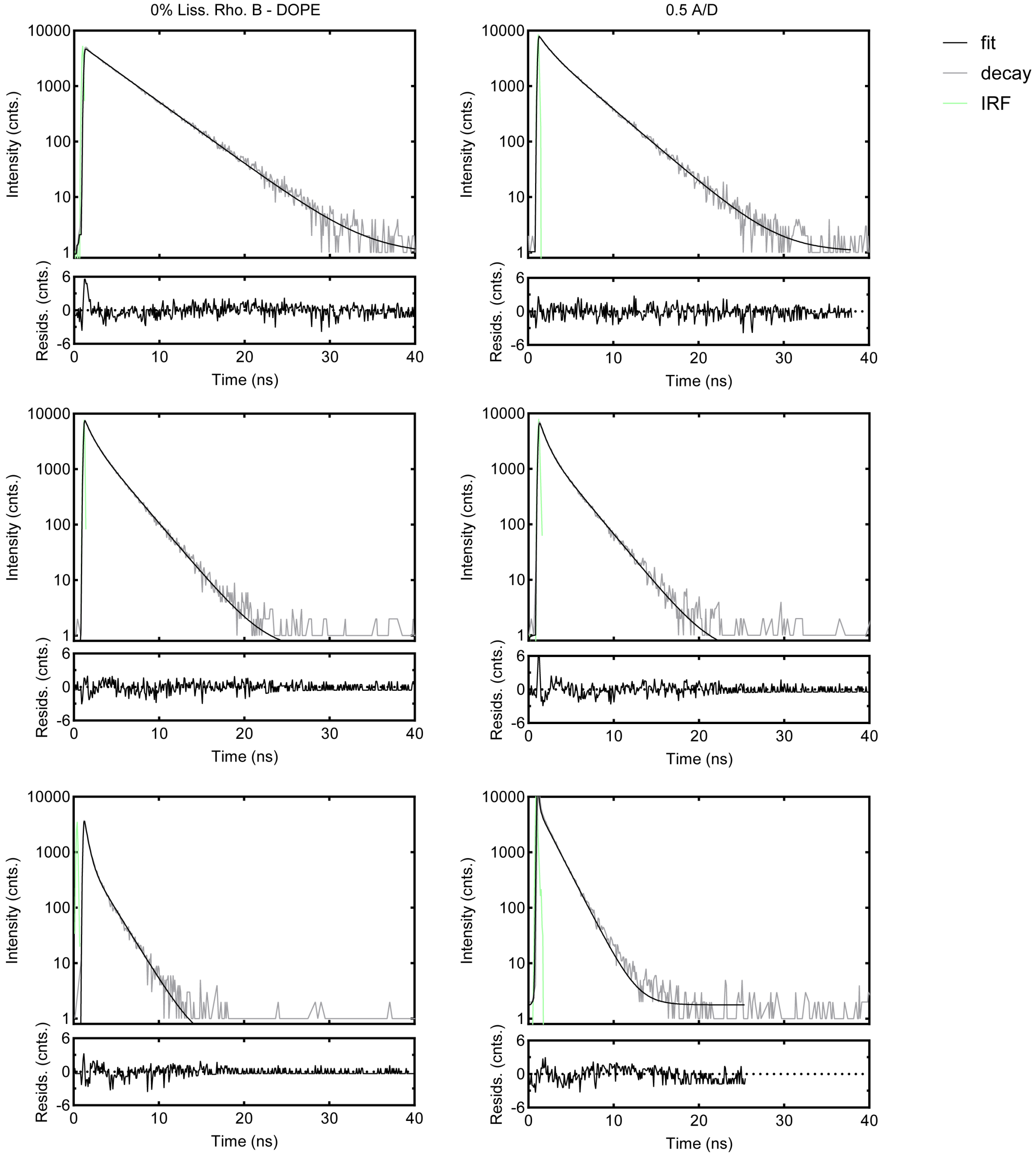


Figure 4: Representative FLIM decay curves from calibration measurements for GUVs containing different acceptor - donor ratios. The donor concentration was kept constant (0.1 mol%), whereas the acceptor concentration was gradually increased ( 0.05%, 0.1%, 0.2%, 0.5%, 1%, and 2%). The decay curve is shown in gray, the fit is shown in black, and the instrument response function (IRF) is shown in green. The concomitant residual counts are plotted below each curve.

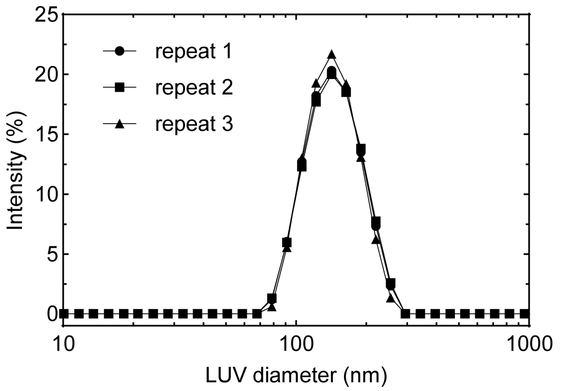


Figure 5: **Determination of the LUV size distribution by dynamic light scattering** For this representative example, the hydrodynamic diameter was 139.5 ± 0.7 nm with a polydispersity index of 0.051 ± 0.023 (n = 3), indicating a monodisperse, unimodal LUV population.

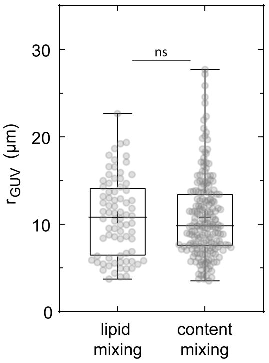


Figure 6: Size distribution of GUVs that were measured in the lipid and content mixing assays. The GUV radius was determined based on confocal images taken at the GUV midplane for the lipid (left; N = 3, n = 78) and content mixing (right; N = 6, n = 200) assays. Statistical significance was tested using a non-parametric Mann Whitney test (ns indicates that there is no significant difference).

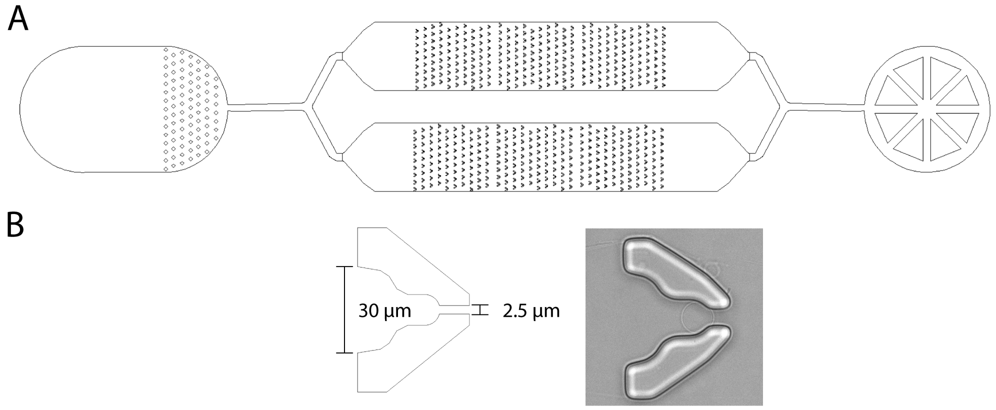


Figure 7: **A.** Design of the microfluidic device for observing LUV-to-GUV fusion in real time. The inlet (left) serves as a reservoir in which the liposome solutions are pipetted. The outlet (right) is connected to a syringe pump via which a negative pressure is applied to draw the liposomes through the device. Liposomes are immobilised in two parallel chambers, each containing 279 traps. **B.** Close-up of a hydrodynamic trap. GUVs enter the structure via a 30 $\mu$m wide opening. Each trap progressively narrows, thereby immobilizing a GUV. We found that a 2.5 $\mu$m wide opening is necessary to create a flow through the trap, otherwise no GUVs are trapped.

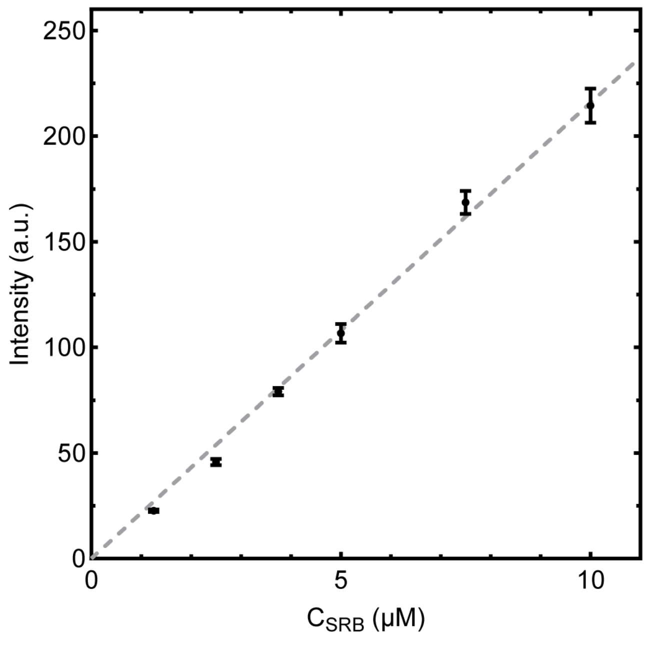


Figure 8: Calibration curve correlating the measured fluorescence intensity signal in confocal images of GUVs containing SRB to the encapsulated SRB concentration. Each point is the average intensity measured in 40 images. Data are represented as mean ± SD. A linear regression was performed with an offset set to zero, yielding y = 21.57 x.

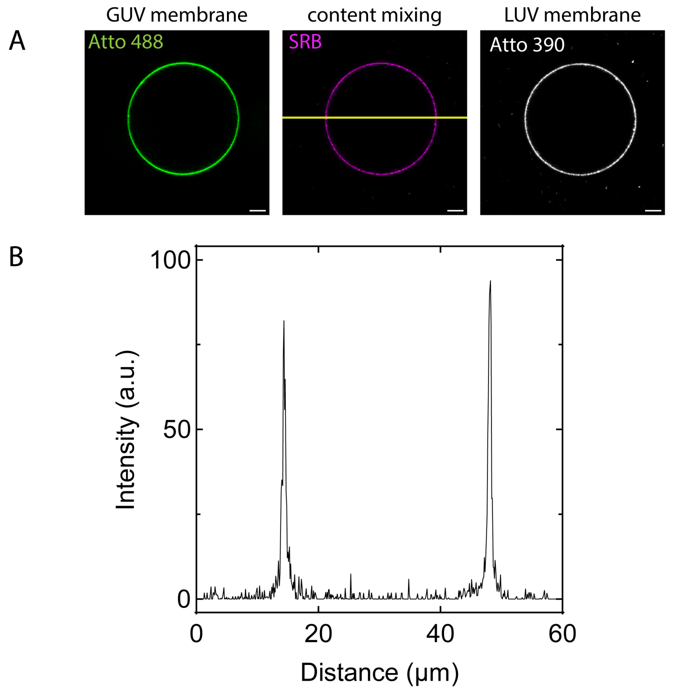


Figure 9: **Control experiment for the content mixing assay. A.** Representative confocal images of a GUV after the docking of LUVs mediated by 105 nM LiNA in bridge mode at 40 °C. The marker for content mixing, SRB, is retained at the GUV membrane, indicating that no transfer of the dye occurs. An intensity profile is acquired along the yellow line. Images are acquired at the equatorial plane of the GUV and scales are 5 micron. **B.** Intensity profile across the GUV in the SRB chanel. There is no increase in SRB intensity observed in the GUV lumen compared to the external medium, indicating that the membrane do not fuse when employing LiNAs in the bridge mode.

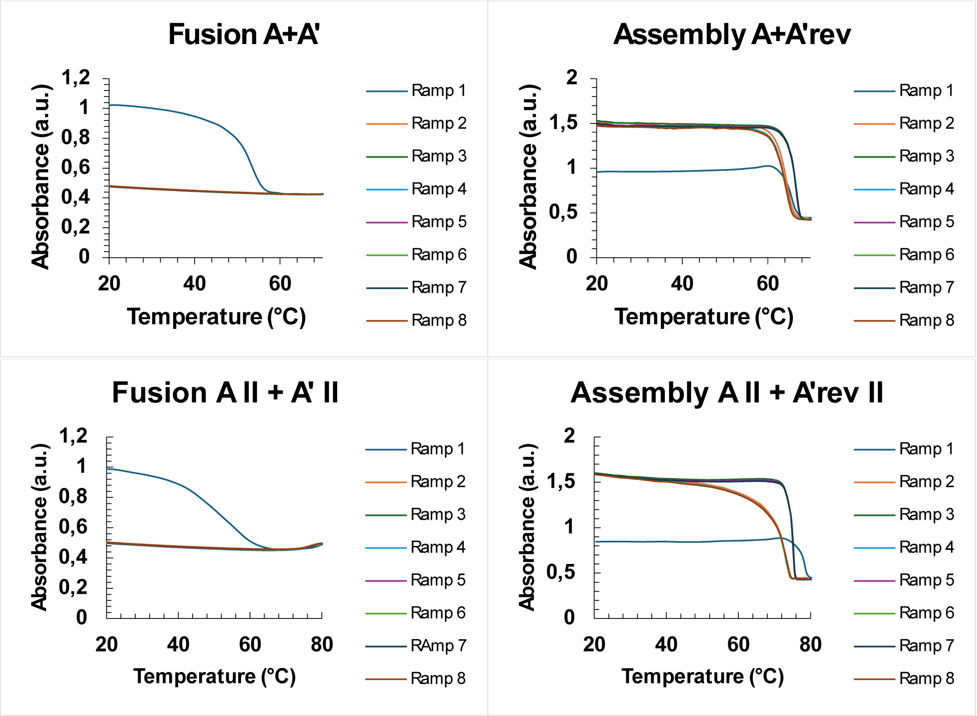


Figure 10: **Determination of Tm for LiNAs in the zipper and bridge mode.** LUVs functionalized with corresponding LiNAs were mixed and the UV absorbance was measured over 8 temperature ramps from 20-70 °C (17mer in zipper mode (A+A') and in bridge mode (A+A'rev)) and 20-80 °C (35mer in zipper mode (AII+A'II) and in bridge mode (AII+A'revII)). For the zipper designs, the first ramp irreversibly fuses the LUVs and no change in absorbance is seen in the following ramps. For the bridge design, the assembly and disassembly of POPC LUVs with LiNA bridges is reversible as the bridge inhibits fusion. For all measurements, ramp 1 is an anealling step with 10 °C/min heating followed by 0.5 °C/min in the following ramps.

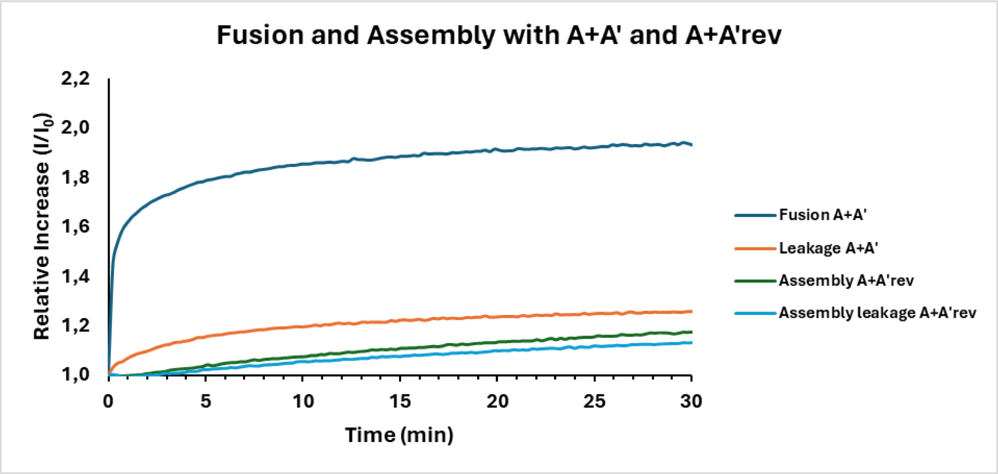


Figure 11: **LiNAs in bridge mode do not result in fusion between distinct populations of LUVs.** An SRB fusion assay was performed for LiNAs in the zipper mode (A+A') and in the bridge mode (A+A'rev) to confirm that the bridge design does not promote fusion. LUVs containing 20 mM SRB were functionalized with 200 nM LiNAs and were mixed with empty LUVs. SRB dilution was observed when employing LiNAs in the zipper mode, but not when using LiNAs in the bridge mode. For leakage experiments, two populations of SRB loaded LUVs were incubated with complementary LiNAs in zipper mode leading to no dilution upon fusion. The same control is performed for leakage upon assembly with LiNAs in bridge mode. The results demonstrate content mixing for the zipper design while the bridge design have a similar increase compared to the passive leakage controls.

Table 1: Oligonucleotide sequences of the LiNA constructs. XN indicates the lipid anchor modification, P3 indicates the triethylene glycol (TEG) spacer. A thymidine residue is added to the anchor-modified terminus to prevent self-aggregation of the amphiphilic LiNAs in aqueous solution. LiNA A and A II were used to functionalize GUVs, whereas other LiNAs were used for functionalizing LUVs.

| LiNA name | Sequence (5'-3') | Mass (calc.; g/mol) | Mass (found, g/mol) | Tm (°C) | $E_{260nm}$ |
|---|---|---|---|---|---|
| A | TXNP3TGTGGAAGAAGTTGGTG | 6476 | 6488 | 54.7 | 182.4 |
| A' | CACCAACTTCTTCCACAP3XNT | 6165 | 6167 | 54.7 | 160.6 |
| A'rev | TXNP3CACCAACTTCTTCCACA | 6165 | 6167 | 54.7 | 162.0 |
| A II | T XNP3TGTGGAAGAAGTTGGTG<br>TGAGAAGTGGTAAAGTAT | 12165 | 12169 | 68.9 | 376.1 |
| A' II | ATACTTTACCACTTCTCACACCA<br>ACTTCTTCCACA **P3XN**T | 11596 | 11597 | 68.9 | 326.1 |
| A'rev II | T XNP3ATACTTTACCACTTCTCA<br>CACCAACTTCTTCCACA | 11596 | 11601 | 68.9 | 326.7 |

Table 2: Membrane compositions of liposomes used in fusion experiments. For each composition, we specify which type of LiNA was used to decorate the liposomes and the corresponding figure panel(s).

| | Liposome | LiNA | Composition | Concentration (mol%) | Figure |
|---|---|---|---|---|---|
| 1 | GUV | A | PC:PE:Ch:Atto488-PE | 49.9:25:25:0.1 | 2B-D, 3, 4, 5 |
| 2 | GUV | A II | PC:PE:Ch:Atto488-PE | 49.9:25:25:0.1 | 2B-D, 4, 5B |
| 3 | GUV | / | PC:PE:Ch:Atto488-PE:Rho-PE | 49.895:25:25:0.1:0.005 | 2E-G |
| 4 | GUV | / | PC:PE:Ch:Atto488-PE:Rho-PE | 49.89:25:25:0.1:0.01 | 2E-G |
| 5 | GUV | / | PC:PE:Ch:Atto488-PE:Rho-PE | 49.88:25:25:0.1:0.02 | 2E-G |
| 6 | GUV | / | PC:PE:Ch:Atto488-PE:Rho-PE | 49.85:25:25:0.1:0.05 | 2E-G |
| 7 | GUV | / | PC:PE:Ch:Atto488-PE:Rho-PE | 49.80:25:25:0.1:0.10 | 2E-G |
| 8 | GUV | / | PC:PE:Ch:Atto488-PE:Rho-PE | 49.70:25:25:0.1:0.20 | 2E-G |
| 9 | GUV | / | PC:PE:Ch:Atto488-PE:Rho-PE | 49.40:25:25:0.1:0.50 | 2E-G |
| 10 | GUV | / | PC:PE:Ch:Atto488-PE:Rho-PE | 48.90:25:25:0.1:1 | 2E-G |
| 11 | GUV | / | PC:PE:Ch:Atto488-PE:Rho-PE | 47.90:25:25:0.1:2 | 2E-G |
| 12 | GUV | A | PC:PE:Ch:PEG2K-PE:Atto488-PE | 49.89:25:25:0.01:0.1 | 6 |
| 13 | LUV | A’ | PC:PE:Ch:Atto488-PE:Rho-PE | 49.40:25:25:0.1:0.50 | 2B-D, 3, 5 |
| 14 | LUV | A’ II | PC:PE:Ch:Atto488-PE:Rho-PE | 49.40:25:25:0.1:0.50 | 5B |
| 15 | LUV | A’rev II | PC:PE:Ch:Atto488-PE:Rho-PE | 49.40:25:25:0.1:0.50 | 2B-D |
| 16 | LUV | A’ | PC:PE:Ch:Atto390-PE | 49.9:25:25:0.1 | 4, 5C, 6 |
| 17 | LUV | A’rev II | PC:PE:Ch:Atto390-PE | 49.9:25:25:0.1 | 4 |
